# Homeostatic cerebellar compensation of age-related changes of vestibulo-ocular reflex adaptation: a computational epidemiology study

**DOI:** 10.1101/2020.08.03.233833

**Authors:** Niceto R. Luque, Francisco Naveros, Eduardo Ros, Angelo Arleo

## Abstract

The vestibulo-ocular reflex (VOR) stabilizes vision during head motion. Age-related changes of vestibular neuroanatomical properties predict a linear decay of VOR function. Nonetheless, human epidemiological data show a stable VOR function across the life span. In this study, we model cerebellum-dependent VOR adaptation to relate structural and functional changes throughout aging. We consider three neurosynaptic factors that may codetermine VOR adaptation during aging: the electrical coupling of inferior olive neurons, the intrinsic plasticity of Purkinje cell (PC) synapses, and long-term spike timing-dependent plasticity at parallel fiber - PC synapses and mossy fiber - medial vestibular nuclei synapses. Our cross-sectional aging analyses suggest that long-term plasticity acts as a global homeostatic mechanism that underpins the stable temporal profile of VOR function. The results also suggest that the intrinsic plasticity of PC synapses operates as a local homeostatic mechanism that further sustains the VOR at older ages. Importantly, the computational epidemiology approach presented in this study allows discrepancies among human cross-sectional studies to be understood in terms of interindividual variability in older individuals. Finally, our longitudinal aging simulations show that the amount of residual fibers coding for the peak and trough of the VOR cycle constitutes a predictive hallmark of VOR trajectories over a lifetime.

## Introduction

Healthy aging progressively degrades postural control and balance (Brandt et al., 2005; Zalewski, 2015; Anson & Jeka, 2016). The consequent loss of static and dynamic balance increases the risk of fall in older adults and hinders their autonomy (Tinetti, 2003; Piirtola & Era, 2006; Desai et al., 2010). Postural control is an adaptive process that relies on the integration of multimodal functions that concurrently mediate body and gaze stability (Mergner & Rosemeier, 1998). In this study, we focus on the vestibulo-ocular reflex (VOR), which ensures gaze stability during head motion (Grossman & Leigh, 1990) by generating rapid contralateral eye movements that stabilize images on the retinal fovea (Fig. 1A). The VOR plays a key role in maintaining balance, and VOR deficits can lead to oscillopsia (i.e., a perturbing illusory oscillation of the visual scene) during locomotion (Demer et al., 1994).

**Figure 1.**
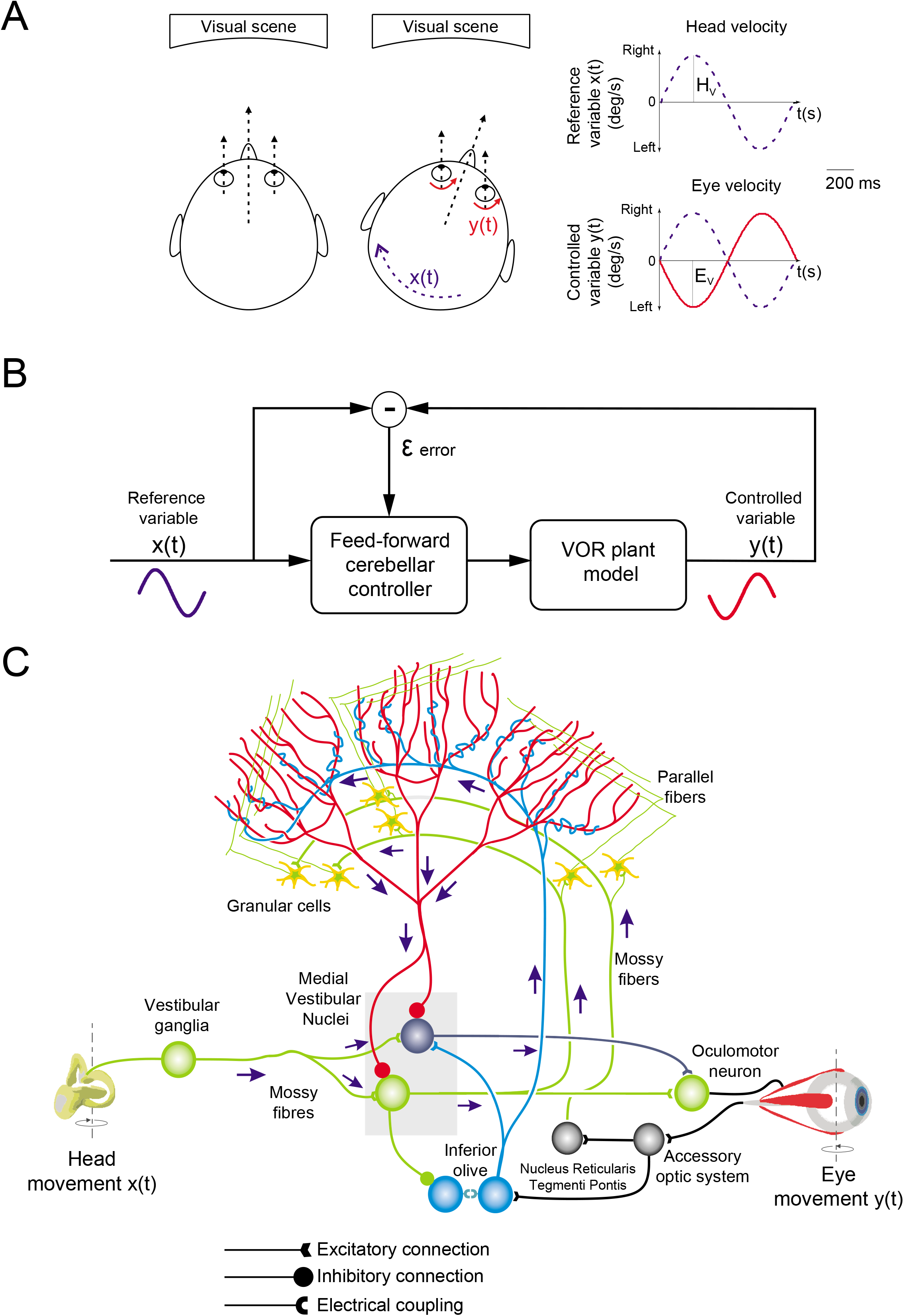
Cerebellum-dependent adaptation of Vestibulo-Ocular Reflex (VOR). **(A)** Horizontal rotational VOR (i.e., r-VOR) stabilizes the visual field during horizontal head rotations, x(t), by producing contralateral eye movements, y(t). **(B)** Cerebellum-dependent VOR adaptation is modelled as a classic feedforward control loop. Cerebellar learning minimizes the error signal *ε*(t), which is computed by comparing the input and output variables, i.e., x(t) and y(t), respectively. **(C)** Schematic representation of the main cerebellar layers, cells, and synaptic connections considered in the model. Mossy fibers (MFs) convey vestibular information onto granular cells (GCs) and medial vestibular nuclei (MVN). GCs, in turn, project onto Purkinje cells (PCs) through parallel fibers (PFs). PCs also receive excitatory inputs from the inferior olivary (IO) system. IO cells are electrically coupled and they deliver an error signal through the climbing fibers (CFs). MVN are inhibited by PCs and they provide the cerebellar output that drives oculomotor neurons. Spike-dependent plasticity occurs at PF-PC and MF-MVN synapses.

The neuroanatomical properties of the vestibular system degenerate with age (Anson & Jeka, 2016; Allen et al., 2017). The number of vestibular receptors (i.e., hair cells) decreases at a rate of about 6% per decade, tending to degenerate from middle age on, independently from pathology. The number of neurons in the vestibular nuclei undergoes a 3% loss per decade, starting at approximately 40 years of age (Lopez et al., 1996; Alvarez et al., 2000). Thus, fewer primary vestibular afferents reach the brain, in particular the downstream areas responsible for VOR adaptation such as the cerebellum (Allen et al., 2017). These structural vestibular losses would predict that aging impairs the detection and encoding of head motion, with a consequent decline of VOR function across the life span. Nevertheless, epidemiological studies report discordant patterns of results regarding VOR functional deficits during normal aging (see Smith, 2016, for a review). Studies in the early 1990s showed age-dependent impairments of the rotatory VOR (r-VOR; Baloh et al., 1993), in response to low-frequency sinusoidal rotations (< 1 Hz; Peterka et al., 1990) as well as to high-amplitude and high-velocity sinusoidal rotations (Paige, 1992). By contrast, recent studies reported that VOR gain (i.e., the ratio between the antagonist eye and head displacements) is preserved even in individuals aged 80 to 90 years. Li et al. (2015) conducted a cross-sectional VOR evaluation on a population of 110 healthy, community-dwelling individuals aged 26 to 92 years. They assessed VOR function with video head-impulse testing (vHIT), which provides a specific clinical assessment of the peripheral vestibular system. They reported that the r-VOR gain remains stable across participants aged up to approximately 80 years and it declines thereafter. McGarvie et al. (2015) measured the VOR gain by using the vHIT for all 6 semi-circular canals across a range of head velocities. They conducted a cross-sectional study on a population of 91 healthy, community-dwelling individuals aged 10 to 89 years (with about 10 subjects per decade). They reported that the r-VOR gain is unaffected by age, because it remained stable even in the group aged 80 to 89 years. Finally, Matiño-Soler et al. (2015) used the vHIT in the lateral semicircular canal plane to evaluate the r-VOR gain as a function of age and head velocity over a population of 212 healthy subjects. They observed a steady r-VOR until 90 years of age for low head impulse velocities and a decline thereafter. At higher-velocity head impulses, they reported a decrease of VOR gain in younger subjects (i.e., from 70 years onwards). Hence, although there are discrepancies in the results, these data consistently support the evidence that VOR function remains unaffected by age until 80-90 years, despite the structural degenerations that impair the vestibular system from middle age on.

Researchers have suggested that some compensatory processes may counter age-related vestibular losses, thus preserving the VOR in older adults (Jahn et al., 2003; Li et al., 2015; McGarvie et al., 2015). However, it is unknown what neural mechanisms are at stake in the brain to maintain VOR function and how they synergistically do so. In this study, we propose a model of cerebellum-dependent VOR adaptation to relate neuroanatomical and functional components of gaze stabilization during head rotations. The goal is to reproduce epidemiological data and to make testable predictions about the interplay of neuronal and plasticity mechanisms operating throughout aging. We hypothesize that three neurosynaptic factors are critical to understand how age affects the VOR:

i. The first neurosynaptic factor is the electrical coupling between inferior olive (IO) neurons through gap junctions (Llinas et al., 1974; Sotelo et al., 1974), which determine the synchronicity and the oscillatory dynamics of the IO network (Lefler et al., 2020). We model the impact of aging on IO electrical coupling by considering the degradation of GABAergic afferents from the medial vestibular nuclei (MVN; Best & Regehr, 2009; Lefler et al., 2014). Because we assume that IO cells encode retina slips during gaze stabilization (Ito, 2013; Luque et al., 2019; Naveros et al., 2019), we assess the consequences of age-related changes in IO electrical coupling on VOR function.
ii. The second neurosynaptic factor is the intrinsic plasticity of Purkinje cell (PC) synapses (Shim et al., 2018; Jang et al., 2020), which modulates the excitability of PCs by adapting their membrane capacitance to morphological changes (Andersen et al., 2003; Zhang et al., 2010). We study the possible role of intrinsic plasticity of PC synapses as a *local* homeostatic process that operates during aging to compensate for the decreasing vestibular inputs as well as for the electro-responsiveness changes in PCs (Andersen et al., 2003; Zhang et al., 2010).
iii. The third neurosynaptic factor is the long-term synaptic plasticity, both potentiation (LTP) and depression (LTD), that drives cerebellar adaptation (Gao et al., 2012; Luque et al., 2019) and sensorimotor learning in general (D’Angelo et al., 2016). We explore the role of LTP and LTD as a *global* homeostatic compensatory process to enhance neural sensitivity during aging.

First, we examine the above neurosynaptic mechanisms independently of one another and we assess their individual impact on VOR adaptation as a function of age. Second, we simulate cross-sectional and longitudinal studies to explore how these mechanisms may codetermine the VOR temporal pattern observed throughout aging. Third, we attempt to identify the factors beneath the large interindividual variability of VOR performance during aging. We test the hypothesis that accounting for the variance in terms of adaptive compensation to residual fibers/connections can help explain the discrepancies between the epidemiological results reported in the literature concerning VOR function in individuals aged 80 to 90 years and thereafter.

## Results

### Cerebellum-dependent VOR adaptation

We framed VOR adaptation within a cerebellum-dependent forward control scheme (Fig. 1B; Lorente de Nó, 1933; Santina et al., 2001; Luque et al., 2019). Computationally, the model reproduces the main properties of the cerebellar circuit and it consists of five neural networks (Fig. 1C). A population of 100 mossy fibers (MFs) conveys primary vestibular inputs (signaling head angular accelerations) onto the cerebellar network. MFs project excitatory afferents onto 200 medial vestibular nuclei (MVN) and 2000 granular cells (GCs). GCs generate a sparse representation of MF inputs and they transmit the encoded sensory information to 200 Purkinje cells (PCs) through excitatory projections. An intrinsic plasticity mechanism (Shim et al., 2018; Jang et al., 2020) regulates the excitability of model PCs, consistently with electrophysiological recordings (Turrigiano et al., 1994; Shim et al., 2017) (see Methods). PCs integrate the afferent signals from PFs (i.e., the axons of GCs), which elicit PC simple spikes (i.e., tonic firing mode, see Luque et al., 2019). PCs also integrate the error-related signal from climbing fibers (CFs), that is inferior olive (IO) cells’ axons. CFs elicit Purkinje complex spikes (i.e., bursting mode). PCs’ responses (either simple or complex spiking) inhibit MVN cells. MVN cells also integrate inputs from MFs and IOs to generate the cerebellar output controlling eye movements (Fig. 1C). The CF-PC-MVN subcircuit comprises two symmetric microcomplexes that control leftward and rightward eye compensatory rotations, respectively (see Methods). Long-term plasticity (LTP and LTD) modulates PF-PC and MF-MVN synapses (Clopath et al., 2014; Badura et al., 2016), whereas the remaining synaptic connections are either non-plastic or electrical, as among IO cells (Fig. 1C; see Methods).

We assessed cerebellum-dependent r-VOR adaptation using a 1 Hz sinusoidal head rotation protocol (i.e., in the natural head rotation range [0.05-5 Hz]; Leigh & Zee, 2015). During 2500 s of simulation (Fig. 2), LTP and LTD plasticity shaped PF-PC and MF-MVN synaptic efficacies (which were randomly initialized) to adapt r-VOR gain and reduce retina slips (i.e., the error sent by IO cells). After about 1000 s, r-VOR gain (averaged over 40 simulated individuals) plateaued at 0.95 (Fig. 2A), a value consistent with experimental VOR data in humans during 1 Hz sinusoidal head rotations (Dits et al., 2013). As encoded by IO cell firing, retina slip errors decreased from 8-9 Hz to 2-3 Hz as VOR accuracy improved (Fig. 2B).

**Figure 2.**
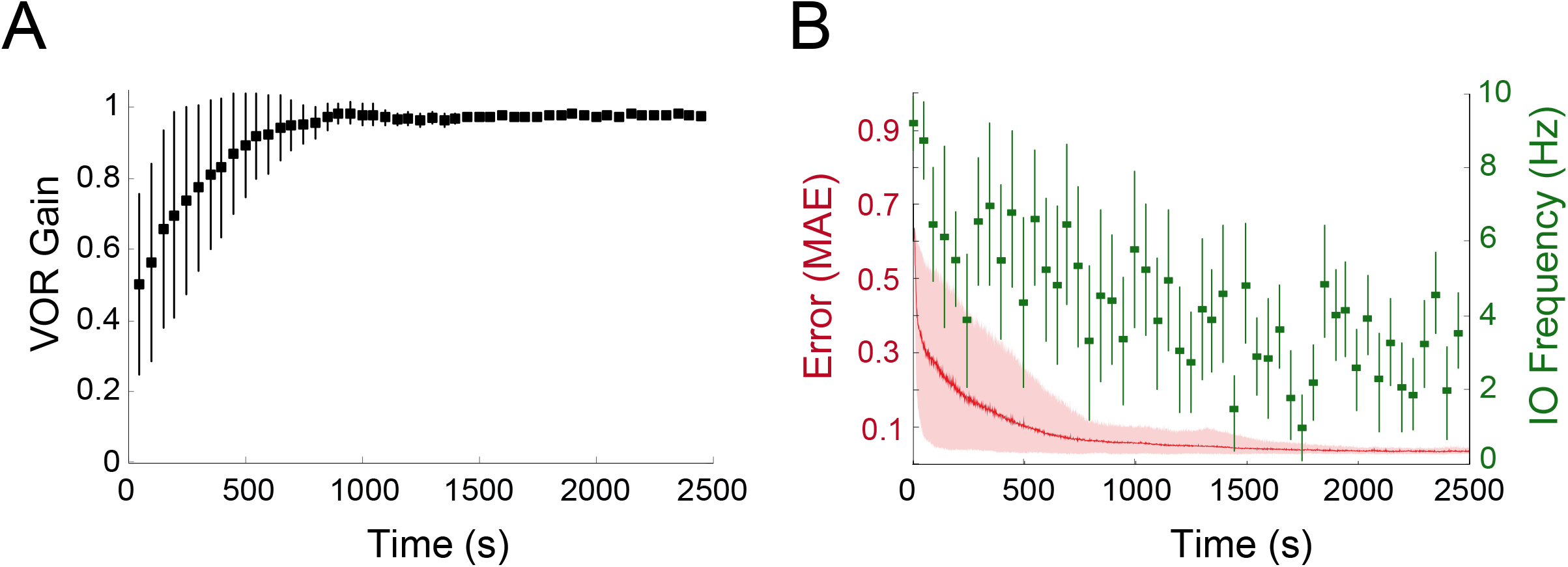
Time course of VOR gain and error during cerebellar adaptation. **(A)** Evolution of the mean VOR gain during 2500 sec of cerebellar learning under a 1-Hz sinusoidal vestibular stimulus (head horizontal rotation). The VOR was averaged across 40 individuals, with each individual obtained by a random initialization of weights at PF-PC and MF-MVN synapses. **(B)** Red curve: mean absolute VOR error during adaptation averaged over 40 individuals. Green squares: mean frequency of IO neurons throughout VOR adaptation.

### Impact of age-related vestibular loss on the electrical coupling of IO neurons

The model IO neurons form an electrically coupled network, whose recurrent dynamics are regulated by the PC-MVN-IO cerebellar loop (Fig. 1C). In particular, the inhibitory MVN action modulates IO electrical coupling strength, which determines the synchronicity of IO firing (Best & Regehr, 2009; Lefler et al., 2014; Najac & Raman, 2015). A strong IO electrical coupling (i.e., a highly synchronized IO network) would allow significant errors to be transmitted to PCs, enabling fast VOR learning (Schweighofer et al., 2013; Tokuda et al., 2013). A reduced IO electrical coupling would lead to slower but more accurate VOR adaptation (e.g., during late learning).

We sought to understand how a progressive age-related decrease of the MVN GABAergic input to IO neurons (owing to vestibular primary afferent loss) would affect the IO network activity (Best & Regehr, 2009; Lefler et al., 2014). We simulated two age groups (20 young subjects: 20 years old; 20 older subjects: 100 years old), and we linearly decreased the inhibitory MVN input to IO as a function of age (from a maximum at 20 years to zero at 100 years; see Methods). We compared the dynamics of IO spatiotemporal firing patterns in a 5×5 lattice configuration (Nobukawa & Nishimura, 2016) after an error-related pulse activated the central IO neuron of the network (e.g., neuron 1 in Supp. Fig. 1A). The electrical coupling between IO neurons produced a rapid transient propagation within the network, eliciting a sequential bursting of IO cells along the lattice’s outward radial direction (Supp. Figs. 1A, B). When comparing the IO network propagation patterns of the two age groups, we found that the central stimulation did elicit more rapid and pronounced membrane potential variations in the IO lattices of older individuals, which resulted in simpler on/off network dynamics as compared to young individuals (Fig. 3A and Supp. Figs. 1B, C). These transient on/off patterns produced a higher mean activation frequency in older IO networks (Fig. 3A). We quantified the complexity of IO spatiotemporal patterns by using the Discrete Wavelet Transform (DWT) (Latorre et al., 2013). We considered these patterns as sequences of images (obtained every ms) and we estimated each image’s the compression by calculating the DWT coefficients. High (low) DWT values corresponded to complex (simple) spatial structures of IO network patterns at a given time. We found that the electrical coupling among IO neurons in older individuals gave rise to significantly simpler spatiotemporal network activations, as compared to young individuals (Fig. 3B; ANOVA F_(294,16)_ =18, p < 10^−7^). This was consistent with a more uniform and synchronized activity of older IO neurons and a higher mean frequency (Fig. 3A). The simpler spatiotemporal dynamics of older IO networks were likely to induce a poorer capacity to encode retina slips. Therefore, we subsequently tested the impact of this less effective error signaling on VOR performance.

**Figure 3.**
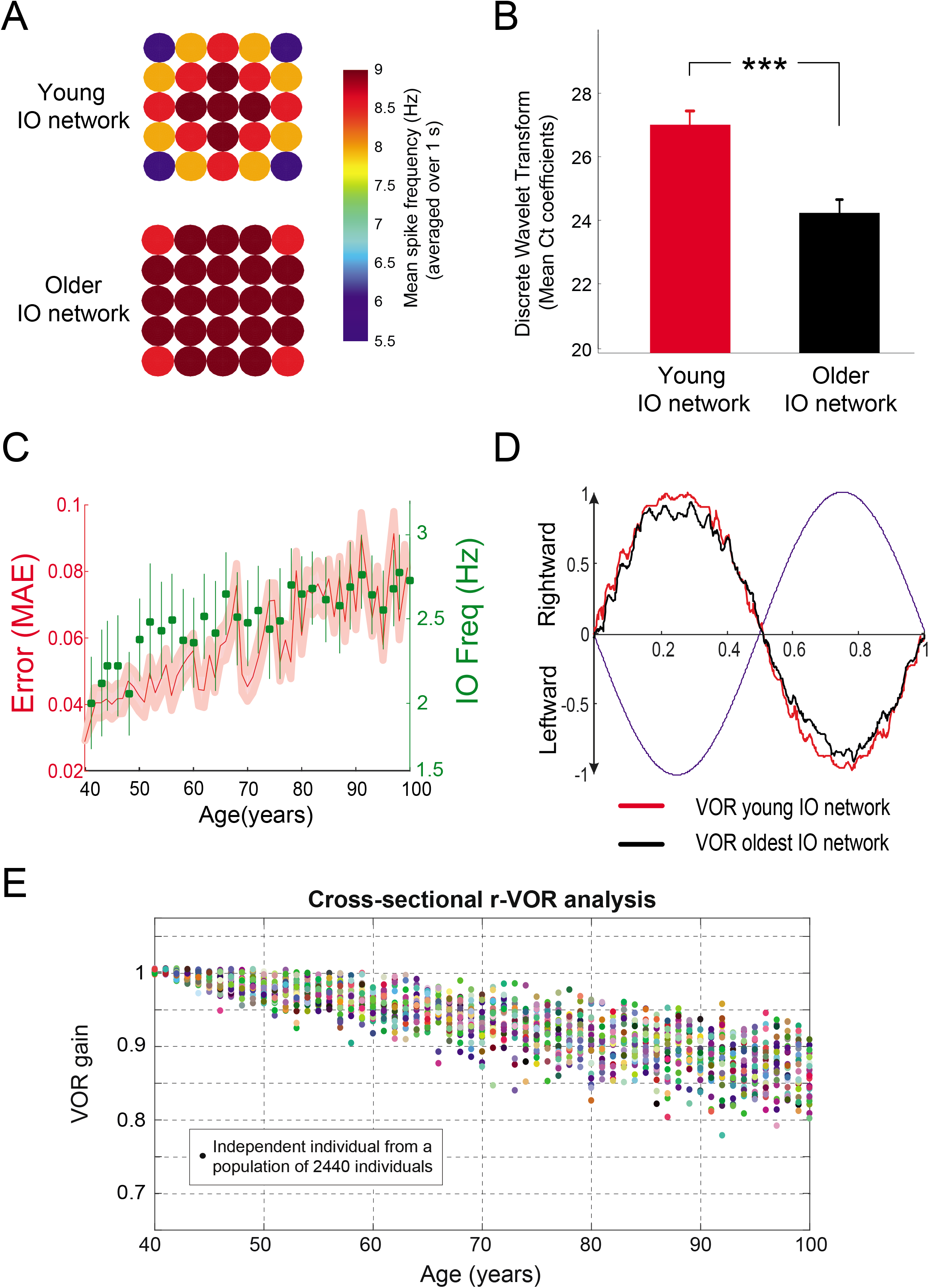
Impact of age-related vestibular loss on IO electrical coupling and cerebellum-dependent VOR adaptation. **(A)** Mean frequency, averaged over 20 young (20 yo, top) and 20 older (100 yo, bottom) individuals, across a cluster of 5 × 5 IO cells (lattice configuration, (Nobukawa & Nishimura, 2016)) during the first second of VOR adaptation under a sinusoidal vestibular stimulus of 1 Hz. **(B)** Discrete Wavelet Transformation (DWT) applied to the snapshot sequence of the IO membrane potentials (Supp. Fig. 1A) obtained during the first second of VOR adaptation for a young (20 yo) and an older (100 yo) individual. **(C)** Mean absolute VOR error (red curve) and corresponding mean IO frequency (green squares), in the presence of altered electrical coupling between IO neurons throughout aging. **(D)** Compensatory eye velocity function in the presence of young (red curve) and older (black curve) IO networks. **(E)** Cross-sectional simulation over a study population of 2440 individuals aged from 40 to 100 years (40 individuals per each year of age). Only the effect of age-related vestibular loss and IO coupling alteration was considered in this simulation, with no compensatory mechanism operating in the downstream cerebellar network. Each individual underwent an independent 1 Hz r-VOR protocol (during 2500 s).

### Impact of age-related vestibular loss on r-VOR performance

We first investigated the consequences of age-related vestibular degradations on VOR function without any compensatory mechanism in the downstream cerebellar network. To do so, we blocked the intrinsic plasticity of PC synapses as well as LTP and LTD at MF-MVN and PF-PC synapses. We simulated a cross-sectional study over a large-scale study population of 2440 individuals aged 40-100 years, by taking a group of 40 individuals per each year of age (i.e., uniform distribution). Each individual underwent an independent 1 Hz head rotation protocol (during 2500 s, as above). At the beginning of the aging simulation, the cerebellar synaptic weights of each 40-year-old individual were obtained at the end of r-VOR learning (Fig. 2). Then, we simulated a loss of primary vestibular afferents as a function of age, based on a degeneration rate of 3% MVN neurons per decade (Lopez et al., 1996; Alvarez et al., 2000). The loss of MVN neurons induced a change in MVN-IO inhibitory action, which gradually increased IO network’s electrical coupling (Best & Regehr, 2009; Lefler et al., 2014; Najac & Raman, 2015). The age-related degradation of primary vestibular afferents also translated into a loss of 0.3% MF-MVN connections per year, as well as 0.3% MF-GC projections per year (starting at 40 years). In addition, the simulated aging process accounted for a loss of approximately 6% of GCs per decade (Bergström, 1973; Baloh et al., 1993; Renovell et al., 2001; Viswasom et al., 2013), which engendered a degradation of 0.6% of PF-PC connections per year. Each of the 2440 individuals independently lost several randomly selected fibers and neurons as a function of age, based on the aforementioned degeneration rates. The aging simulation results showed a steady decline of VOR function (Figs. 3C-E), with the accuracy of r-VOR gain significantly impaired with aging. Across the study population, VOR gain declined quasi-linearly as a function of age (−0.25 %/year; Fig. 3E).

### Intrinsic plasticity at PC synapses as a local homeostatic mechanism

The detailed PC model reproduced the three characteristic spiking patterns observed experimentally (Fig. 4A): simple spiking response (i.e., tonic firing at 10-250 Hz), complex spiking response (i.e., bursting activity up to 600 Hz), and post-complex spike pauses. We previously showed that PC spike burst-pause dynamics are likely to play a key role in VOR adaptation and reversal learning (Luque et al., 2019). Here, we investigated the consequences of age-dependent changes of PC excitability on VOR function. With aging, the number and the surface of PC synapses decrease significantly (Zhang et al., 2010). We reasoned that intrinsic plasticity could adapt PCs’ response during aging, thus operating as a local homeostatic mechanism. The membrane capacitance of modeled PCs decreased as a function of age (Li & Li, 2013) (Fig. 4B; see Methods). This led to increased tonic firing rates in older PCs (Fig. 4C), consistent with electrophysiological data (Zhang et al., 2010). We also assessed the relationship between the duration of post-complex spike pauses and the duration of pre-complex spike ISIs in model PCs. We realized this measure by incrementally increasing PF inputs while maintaining the CF stimulation constant (i.e., only ISIs immediately following complex spikes were considered for this analysis, as in experimental data by Grasselli et al., 2016). The PC model with the intrinsic plasticity mechanism predicts that the linear relation between the duration of post-complex spike pauses and the duration of pre-complex spike ISIs would be preserved during aging (Fig. 4D; R^2^ = 0.9932; p < 10^−4^).

**Figure 4.**
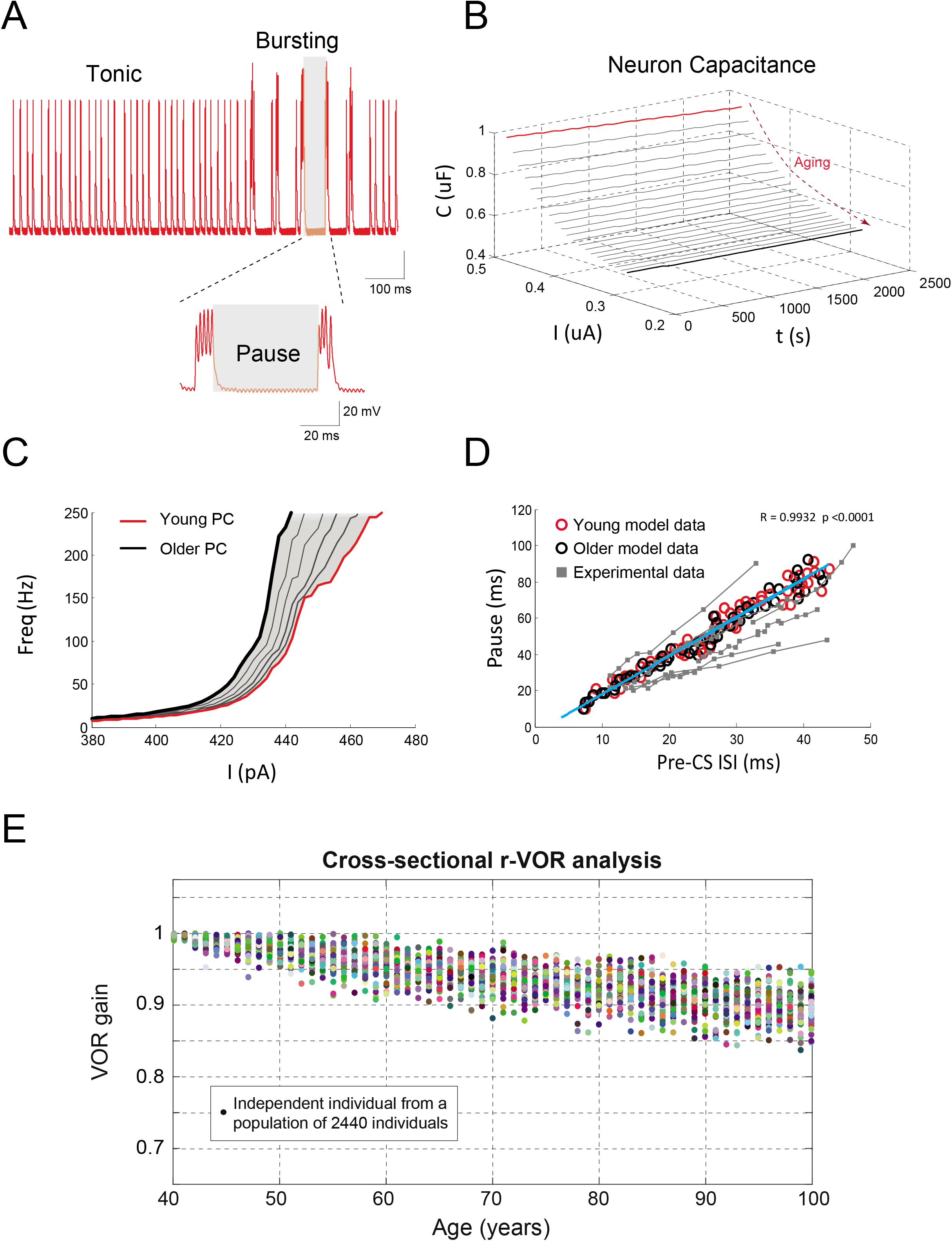
Intrinsic plasticity of PC synapses acts as a local homeostatic mechanism. **(A)** Trimodal spiking patterns of modeled PCs: tonic firing, corresponding to simple spikes elicited by PF inputs; bursting mode, during which complex spikes (bursts of spikes) are elicited by CFs (∼500 synapses, (Palay & Chan-Palay, 2012)) that can even suppress simple spiking; and silent mode, corresponding to an extended hyperpolarization period called ‘post-complex spike pause’. **(B)** Modulation of the membrane capacitance of modelled PCs as a function of age by means of intrinsic plasticity. **(C)** Firing rate of PCs in young vs. older individuals measured with PCs operating in spiking tonic mode (10-250Hz). **(D)** Correlation between pause duration and pre-complex spike ISI duration in modelled young and older PCs (red and black circles, respectively), as well as in real PCs (gray squares, (Grasselli et al., 2016)). **(E)** Cross-sectional simulation over a study population of 2440 individuals (aged from 40 to 100 years, with 40 individuals per each year of age) accounting for vestibular loss, IO electrical coupling chances, and intrinsic plasticity of PC synapses.

We then ran a second aging simulation to test to what extent PC intrinsic plasticity may act when the vestibular loss occurred. We simulated a cross-sectional study by taking a sample of 2440 individuals (age range: 40-100 years; 40 individuals for each year of age). Each individual underwent the same VOR adaptation protocol (1 Hz sinusoidal rotation during 2500 s). The initial conditions (in terms of cerebellar synaptic weights) corresponded to those obtained after r-VOR learning per each independent individual (Fig. 2A). Age-dependent structural degenerations translated into a loss of 0.3% and 0.6% MFs and PFs per year, respectively. As a consequence of MVN loss, IO electrical coupling progressively increased with age (Best & Regehr, 2009; Lefler et al., 2014; Najac & Raman, 2015). LTP and LTD plasticity at MF-MVN and PF-PC synapses were blocked to isolate the effect of the local homeostatic mechanism provided by the intrinsic plasticity of PCs. We found that PCs’ increasing excitability could only partially counterbalance the decreased depolarizing currents elicited by PFs throughout aging. This resulted in a quasi-linear decrease of r-VOR gain across lifetime (−0.17 %/year), along with an increasing interindividual variability (Fig. 4E).

### Cerebellar spike-timing dependent plasticity as a global homeostatic compensatory mechanism

We then tested whether LTP and LTD at PF-PC and MF-MVN synapses could enhance the sensitivity of PCs and MVN to degraded inputs during aging. First, we analyzed the weight distributions at PF-PC and MF-MVN synapses after r-VOR learning as a function of age. We compared the synaptic weights of simulated young and older individuals (20 and 100 years, respectively). In both age groups, cerebellar learning led to antisymmetric weight distributions at both PF-PC and MF-MVN synapses (Fig. 5), corresponding to the two microcomplexes that controlled rightward and leftward eye movements. As expected, PCs’ inhibitory action onto MVN generated opposite weight patterns at PF-PC compared to MF-MVN synapses (Figs. 5A, B vs. 5C, D). In older individuals, an increase the remaining fibers’ weights compensated for the loss of vestibular afferents (Figs. 5B, D). When comparing the distributions obtained by the normalized sums of synaptic weights across PFs (i.e., to estimate the input drive received by PCs), we found platykurtic-like distributions in older individuals (Figs. 5B, D) as compared to more leptokurtic profiles in young individuals (Figs. 5A, C). The ratio between the number of saturated synaptic weights and the number of active afferents increased with age: 28% in young vs. 64% in older PF-PC synapses (Fig. 5A vs. 5B); and 21% in young vs. 31% in older MF-MVN synapses (Fig. 5C vs. 5D). Consequently, the neural drive (defined as the area obtained by convolving a unitary impulse signal with the weight distributions) increased with age: it was 2.64 times larger in the older PF-PC synaptic distribution and 1.64 times larger in the older MF-MVN synaptic distribution, as compared to younger individuals, respectively.

**Figure 5.**
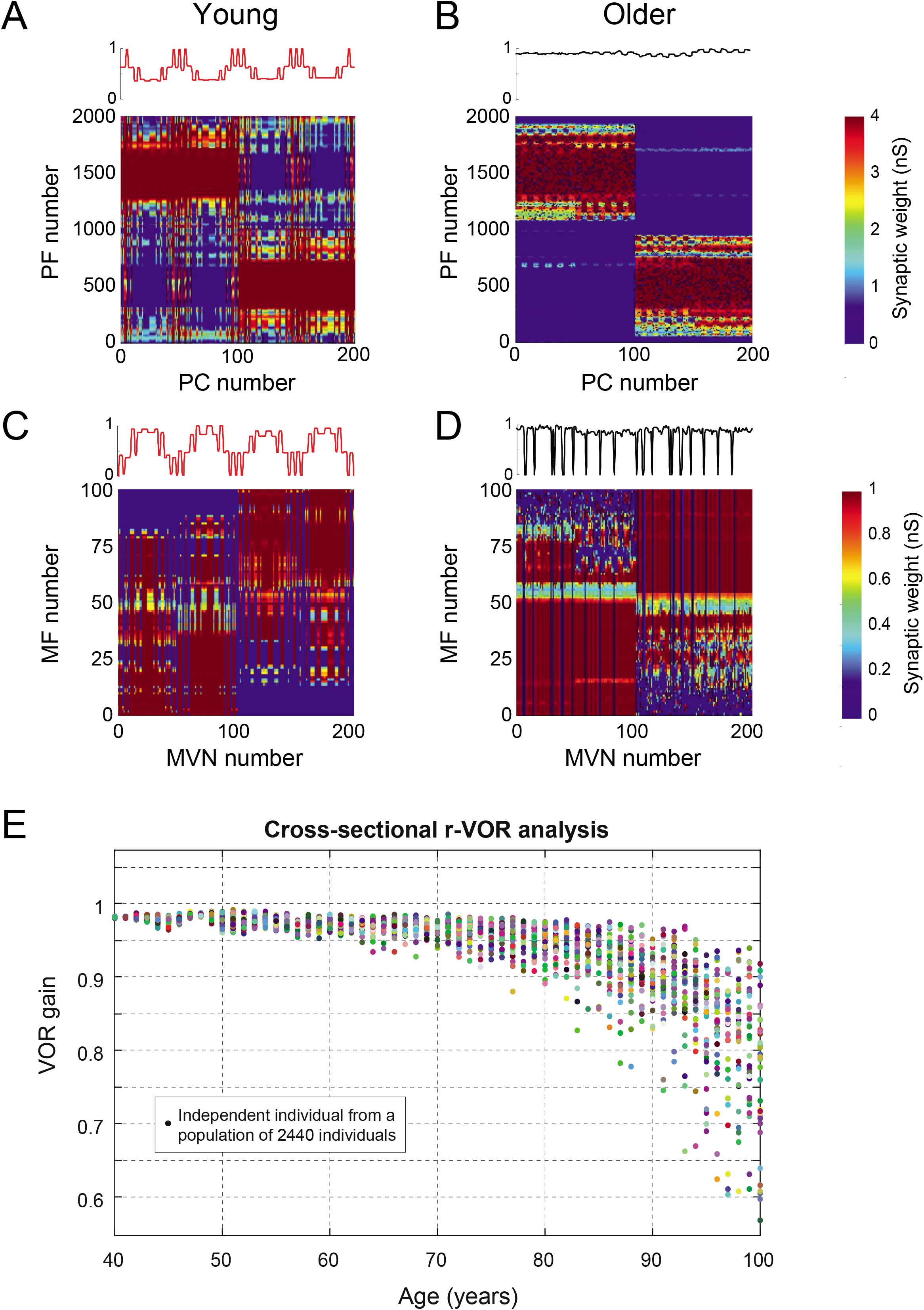
Cerebellar LTP/LTD operates as a global homeostatic compensatory mechanism. **(A, B)** Synaptic weight distributions obtained at PF-PC connections by averaging over 20 young individuals (20 yo) and 20 older ones (100 yo). Each individual underwent an independent 1 Hz r-VOR adaptation (during 2500 s). The weights that correspond to the two cerebellar microcomplexes devoted to the control of rightward and leftward eye movements are visible. **(C, D)** Synaptic weight distributions at MF-MVN connections by averaging over 20 young individuals (20 yo) and 20 older ones (100 yo). The antisymmetric distributions with respect to (A, B) are caused by the inhibitory PC projections onto MVN. **(E)** Cross-sectional aging simulation accounting for vestibular loss, IO electrical coupling alterations, and LTP/LTD at PF-PC and MF-MVN synapses.

We then ran a third cross-sectional aging simulation to isolate the role of LTP and LTD in preserving VOR across a lifetime (i.e., by blocking intrinsic plasticity of PC synapses). We considered another study population of 2440 individuals (age range: 40-100 years; 40 individuals per year of age), and we applied the same VOR adaptation protocol (1 Hz head rotation during 2500 s). We found that compensatory LTP and LTD at PF-PC synapses and MVN synapses successfully preserved sensorimotor associations underpinning VOR until approximately 80 years of age (Fig. 5E). After that, we observed increasing performance variance across individuals, and r-VOR gain declined in the group aged 85-100 years (−0.9 %/year).

### Impact of aging on the VOR function: Full cross-sectional study

We then combined all age-related factors and compensatory mechanisms examined so far to assess their synergistic impact on r-VOR adaptation. We ran a fourth cross-sectional analysis on another cohort of 2440 individuals (age range: 40 - 100 years), with each individual undertaking the same r-VOR adaptation protocol (1 Hz sinusoidal head rotation during 2500 s). Intrinsic plasticity of PC synapses and LTP and LTD at MF-MVN and PF-PC synapses provided local and global homeostatic adaptation, respectively. The computational epidemiological results suggested that the considered neurosynaptic factors concurrently shaped r-VOR function across the life span. VOR gain remained unaffected by age until approximately 80-90 years (Fig. 6A). PCs’ intrinsic plasticity further sustained VOR function compared to when only LTP and LTD were present (Fig. 5E). The variability across individuals increased significantly after 90 years (Supp. Fig. 2), and VOR gain declined steadily thereafter (−0.8 %/year).

**Figure 6.**
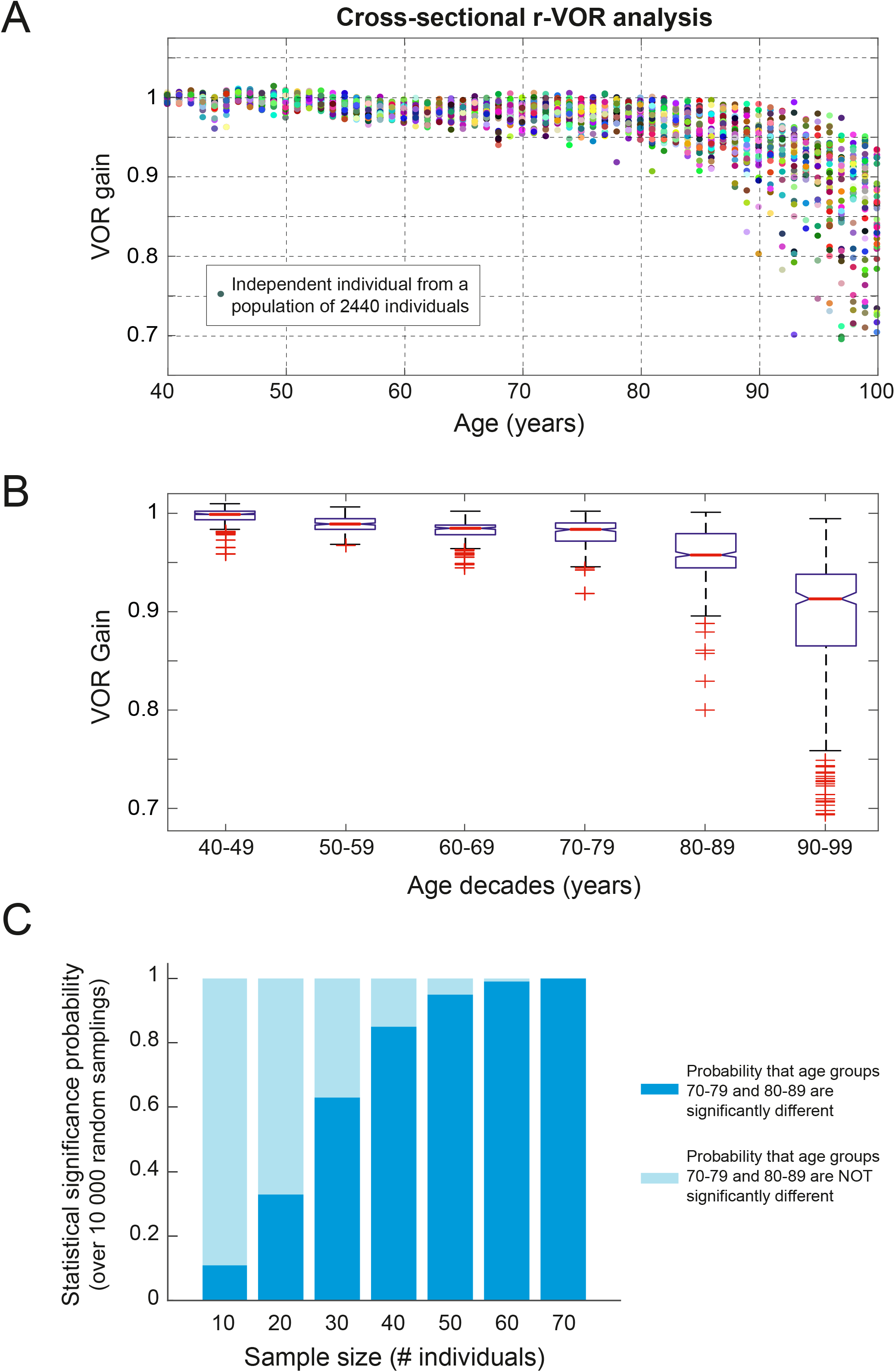
Impact of aging on VOR: cross-sectional study. **(A)** Cross-sectional aging simulation accounting for vestibular loss, IO electrical coupling changes, PC intrinsic plasticity, and LTP/LTD at PF-PC and MF-MVN synapses. A study population of 2440 individuals (age range: 40-100 years; 40 individuals per each year of age) underwent the same r-VOR protocol (1 Hz head rotation during 2500 s). **(B)** Comparison of r-VOR gain across decade age bands (with 400 individuals per decade). **(C)** Probability that r-VOR gain in age groups 70-79 and 80-89 is statistically different as a function of sampling size (computed over 10 000 random iterations per each sampling size).

To compare our results against human cross-sectional data (Li et al., 2015; Matiño-Soler et al., 2015; McGarvie et al., 2015), we analyzed VOR gain changes across decade age bands (Fig. 6B). Each age band included 400 simulated individuals. A multiple comparison post hoc test showed that VOR function began to decline during the 80-89 age band, and it continued to decrease thereafter (ANOVA F_(7,3032)_=736.2, p < 10^−9^). Then, given the large interindividual variability observed at older ages (Figs. 6A, B), we investigated to what extent the sample size may affect the outcome of multiple comparison analyses across age groups. We considered a set of different sample sizes: 10, 20, 30, etc. For each sample size (e.g., 10), we randomly sampled a corresponding number of individuals (e.g., 10) from the population of 400 individuals per each decade age band. Then, we ran a multiple comparison post hoc analysis across age groups. We repeated this process 10000 times and we computed the overall probability of observing a statistically significant difference between VOR gains (Fig. 6C). This model-based analysis showed that a sample size of 50, or greater, individuals per decade would capture the statistical difference between the 70-79 and 80-89 age groups with a probability of 0.95. It also suggested that: for a sample size of 10 individuals per decade (as in McGarvie et al., 2015), the probability of observing a significant r-VOR difference between 70-79 and 80-89 age groups would be as low as 0.11; for a sample size of 20 individuals per decade (Matiño-Soler et al., 2015), the predicted statistical significance probability would be as low as 0.33; and for a sample size equal to 30 individuals per decade (Li et al., 2015), the probability of observing a significant statistical difference would be equal to 0.63.

### Impact of aging on the VOR function: Full longitudinal study

We ran a longitudinal aging simulation by combining all considered age-related factors and compensatory mechanisms. We took a study population of 40 individuals, and we simulated a 61-year follow-up for each individual (i.e., from 40 to 100 years). Again, we considered a loss of 0.3% and 0.6% of MFs and PFs loss per year, respectively. Also, age-related changes of MVN GABAergic inputs to IO shaped the electrical coupling within IO network as before. However, in contrast to previous cross-sectional scenarios, these neural losses accumulated across each individual’s lifetime. VOR performance across the study population remained unchanged by age until approximately 85-90 years, and it declined afterward (Fig. 7A). The interindividual variability increased significantly during the last 15-year period (85-100 years), becoming approximately five times larger at 100 years as compared to 85 years (Supp. Fig. 2).

**Figure 7.**
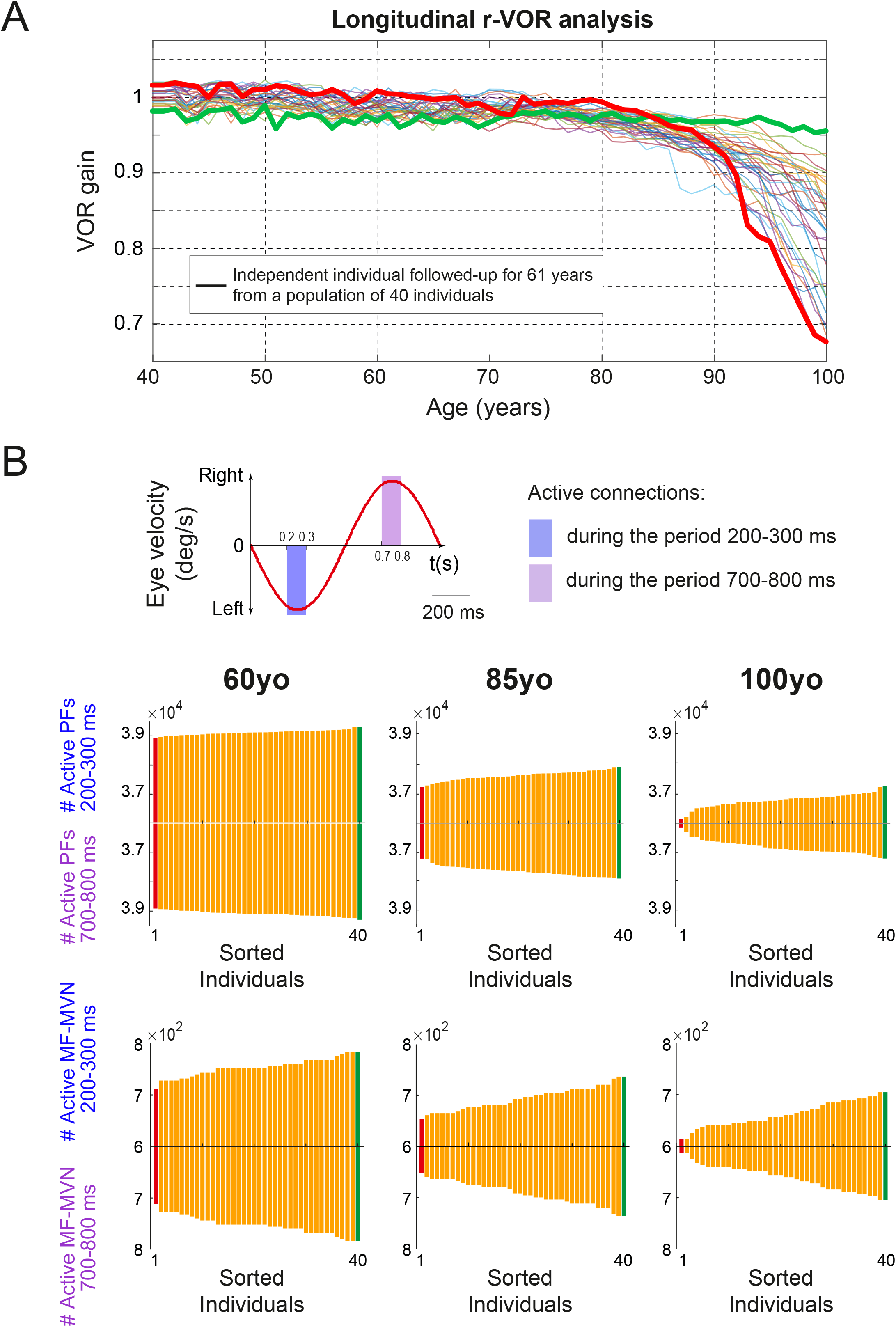
Impact of aging on VOR: longitudinal study. **(A)** Longitudinal aging simulation accounting for vestibular loss, IO electrical coupling changes, PC intrinsic plasticity, and LTP/LTD at PF-PC and MF-MVN synapses. A study population of 40 individuals underwent a follow-up evaluation during 60 years (from 40 to 100 years of age), i.e., age-related changes accumulated longitudinally for each individual. The thick green and red curves correspond to the two individuals with best and worst VOR performance at 100 years of age, respectively. **(B)** Top: 1-Hz eye velocity function. The colored time windows correspond to the peak and the trough of the sinusoidal profile (i.e., between 200-300 ms and 700-800 ms). Center & Bottom: Residual PFs (center) and MF-MVN projections (bottom) active at the peak and the trough of the eye velocity profile across the study population. Individuals are sorted in ascending order on the basis of their VOR performance. The red and green individuals correspond to the worst and best 100-year-old individuals in A, respectively. Left column: distribution of residual PFs and MF-MVN connections at 60 yo. Central column: at 85 yo. Right column: at 100 yo.

Finally, we investigated the underlying factors that determined the difference between steady and declining VOR trajectories in individuals aged 85-100 years (e.g., thick green curve vs. thick red curve in Fig. 7A, respectively). Knowing that: (i) all individuals of the same age had the same probability of losing primary vestibular afferents, MVN, MFs, and PFs; (ii) the degeneration process affected fibers and neurons based on random selection; (iii) the local and global homeostatic mechanisms (i.e., intrinsic plasticity and LTP and LTD) operated equally across all individuals, we reasoned that a possible determinant of VOR aging trajectory could lie in the distribution of the remaining fibers/synaptic connections post-age-related loss. We hypothesized that the activity of some subsets of the remaining connections might be more critical than others in terms of information content for the encoding of sensorimotor associations and then for maintaining VOR function. To test this hypothesis, we first sorted all individuals based on their VOR gain performance at 100 years. Then, we compared the subsets of residual connections active at specific VOR cycle moments across individuals. We found that the number of remaining connections responsible for the encoding of the peak and the trough of the eye velocity function correlated significantly with VOR performances of 100-year-old individuals (Fig. 7B and Supp. Fig. 3). That is, the sorting of 100-year-old individuals based on their residual VOR performance matched the sorting of the same individuals based on the number of residual PFs and MF-MVN projections that coded for the sinusoid’s peak and trough. This correlation held already at 85 years of age (i.e., the cut off age between steady and declining VOR trajectories; Fig. 7B) and even at 60 years of age (i.e., numerous years before the discontinuity-time point between “good” and “bad” aging trajectories; Fig. 7B). Given that the overall number of lost connections was the same across all the study population, this implied that those individuals that *by chance* had most of the remaining connections involved in the encoding of those two critical moments in the VOR period (i.e., between 200-300 ms and 700-800 ms) had the best chances to preserve their VOR performances throughout aging.

## Discussion

This study presented a computational epidemiological model of cerebellum-dependent VOR adaptation. The proposed simulation framework attempted to provide a mechanistic insight into the factors that determine the impact of aging on rotatory VOR function. The model captured the neural computations at stake in the cerebellar circuit and it reproduced the biophysical properties of Purkinje cells (PCs). Importantly, the computational cross-sectional and longitudinal analyses presented in this work allowed the discrepancies among human VOR aging studies in the literature to be understood in terms of interindividual variability (in particular, in individuals over 80 years of age).

We tested the hypothesis that three neurosynaptic factors are key to relate age-related structural and functional VOR changes: the electrical coupling of inferior olive (IO) neurons, the intrinsic plasticity of PC synapses, and LTP and LTD at PF-PC as well as MF-MVN synapses. To verify this hypothesis, we first ran a series of simulations to isolate the role of these three factors in determining VOR changes throughout aging. We found that vestibular structural losses caused by aging have consequences on the spatiotemporal activity patterns of the IO network. IO neural activity becomes simpler, similar to an on/off ensemble dynamic. This on/off network dynamic reduces the accuracy of retina slip coding, which in turn impairs VOR adaptation. We tested the consequences on r-VOR aging by running a cross-sectional epidemiological simulation that isolated the effect of vestibular loss and IO coupling alteration, while blocking all compensatory mechanisms in the downstream cerebellar network. As expected, we found a linear decline of r-VOR gain as a function of age, determined by steady vestibular losses. This result contrasts with human epidemiological data, which show a well-preserved VOR function even in individuals aged 80 to 90 years (Li et al., 2015; Matiño-Soler et al., 2015; McGarvie et al., 2015). We then ran another cross-sectional simulation to assess the local homeostatic action of PCs’ intrinsic plasticity, which adaptively increases PCs’ neural excitability to signals transmitted by cerebellar granule cells. At the level of single PCs, we found that tonic firing rates increased with age, according to experimental data (Zhang et al., 2010), whereas the linear relation between the duration of post-complex spike pauses and the duration of pre-complex spike ISIs remained unchanged (testable prediction). At the level of VOR function, we found that intrinsic plasticity of PC synapses could moderately counter the vestibular structural losses, resulting in a slower, but still linear decline of VOR accuracy over time. This result again contrasts with human epidemiological data. We then investigated the impact of LTP and LTD at PF - PC and MF - MVN synapses to study a global homeostatic process to adapt cerebellar synaptic weights to degrading vestibular inputs.

Our third cross-sectional aging simulation indicated that LTP and LTD can sustain VOR function by enhancing the neural sensitivity to residual afferent signals throughout aging (i.e., it allowed the full synaptic range to be exploited in order to preserve the neuronal drives). However, the compensatory action by LTP and LTD became ineffective in the presence of significant levels of vestibular losses (i.e., beyond 85 years) because of synaptic weight saturation. This result is consistent with the saturation hypothesis of Nguyen-Vu et al. (2017). They showed that a specific type of pre-training that desaturates synapses can improve the ability of mutant mice to learn an eye movement task. Conversely, they found that a specific procedure that saturates synapses can impair the learning ability. In our model, the progressive saturation of PF-PC and MF-MVN synapses limited VOR adaptation, thus impairing the compensatory action of LTP and LTD in the oldest individuals.

We then ran a fourth cross-sectional simulation to assess how the three neuro-synaptic factors would work concurrently during VOR aging. We found that the global homeostatic compensation mediated by cerebellar LTP and LTD was indeed primarily responsible for preserving VOR gain. The results also showed that the local homeostasis implemented by the intrinsic plasticity of PC synapses played a role in further sustaining VOR gain in individuals 80-90 years old and in attenuating its decline afterwards. The slope attenuation was within the same range as the one recently reported in a study of PCs’ intrinsic plasticity during long-term VOR consolidation in mice (Jang et al., 2020). In addition, the computational epidemiological results allowed us to evaluate the possible role of interindividual variability in biasing human VOR aging analyses, thus leading to discordant conclusions among previous studies in the literature (Smith, 2016). The model predicted that a sample size of 10 and 20 individuals per decade, as done in Matiño-Soler et al. (2015) and McGarvie et al. (2015), respectively, would led to very low probabilities of observing statistically significant VOR differences in those aged 70-79 versus those aged 80-89 (0.11 and 0.33, respectively). Accordingly, both Matiño-Soler et al. (2015) and McGarvie et al. (2015) reported a well-preserved VOR function across the life span (at least until 90 years of age). Our model also suggested that a sample size of 30 individual per decade, as used by Li et al. (2015), would lead to a probability of 0.63 of observing a VOR accuracy drop between the 70-79 and 80-89 age groups. Accordingly, and in contrast to other epidemiological studies, Li et al. (2015) reported some evidence for a decline of VOR function after 80-85 years of age.

Finally, we further exploited the presented computational epidemiology model to run a longitudinal aging simulation. This allowed us to follow individual aging trajectories over 60 years in an attempt to better understand the factors determining interindividual differences across aging (i.e., differentiating steady and declining VOR trajectories). Importantly, we found that the number of remaining PFs and MF-MVN projections coding for the peak and the trough of the VOR cycle provided a predictive hallmark for the VOR aging trajectory on a single-subject basis. That is, individuals who lacked active PF and MF-MVN afferents in those precise moments were robustly predicted to have more difficulties in VOR adaptation throughout aging. This prediction likely could be tested in animal models by seeking those fibers that are most active when the ocular velocity is maximal during VOR. For example, the identification of specific cerebellar GCs (and of their PFs) that are active upon induction of specific stimulation is possible during in vivo mice experiments (Ishikawa et al., 2015).

The cerebellar model presented here made several assumptions. The first assumption was that the GC layer univocally encoded vestibular (related to head motion) signals through the temporal activation of non-overlapping cell populations during cerebellar VOR adaptation. GCs are thought to encode vestibular signals into sparse representations allowing interferences across tasks to be minimized and neuronal resources to be optimized by reducing redundancy (D’Angelo & De Zeeuw, 2009). The recurrent inhibitory Golgi cells - GC connections suggest the granular layer may act as a recurrent dynamic network (Yamazaki & Tanaka, 2005). Thus, GCs are likely to generate a randomly repetitive network response characterized by active/inactive state transitions with no repetition of active cell populations (Yamazaki & Tanaka, 2007). The model also assumed a progressive degradation of vestibular afferents integrated by the GC layer with aging (Bergström, 1973; Baloh et al., 1989), which led to a degradation of PFs, which in turn impaired long-term PF-PC synaptic adaptation. Notably, neural regeneration can occur at PF-PC synapses thanks to the Glu*δ*2 receptor (Ichikawa et al., 2016), whereas Glu*δ*2 deficits lead to disruption of LTD at PC synapses and motor impairment in VOR tasks (Yuzaki, 2013; Pernice et al., 2019). Some evidence has suggested that neural loss can be related to the absence of the Glu*δ*2 receptor, because the deletion of GluR*δ*2 expression in mutant mice (GluR*δ*2ho/ho) induces PC and GC reduction over a lifetime (Zanjani et al., 2016). A gradual decrease of Glu*δ*2 with aging would compromise Glu*δ*2-dependent processes that would then reduce intrinsic PC excitability and eventually impair LTD at PCs.

The model also assumed a compromised IO electrical coupling due to degraded GABAergic afferents from MVN during aging. The strength of gap junctions among olivary neurons was modelled as asymmetric (De Zeeuw et al., 1998; Lefler et al., 2014). The level and the direction of this asymmetry were regulated by emulating the GABAergic feedback (Lefler et al., 2014). The coupling asymmetry allowed for the creation of different spatial configurations of PCs’ complex spike patterns. The GABAergic inputs from MVN could directly cause a transient decrement in electrical coupling among IO cells (Lefler et al., 2014). GABAergic feedback not only temporarily blocked the transmission of signals through the olivary system but it also could isolate IO neurons from the network by shunting the junction current (Loewenstein, 2002). In the absence of GABAergic feedback, electrical coupling was not counteracted, and IO network oscillations were not mitigated but rather increased. There is only indirect evidence for an age-related degeneration of the GABAergic MVN inputs throughout aging. The r–aminobutyric acid, GABA, inhibits the formation of lipoxidation end products (Deng et al., 2010). The presence and accumulation of lipofuscin with aging, a lipoxidation product, are essential parts of the traditional theory of aging (Sulzer et al., 2008). Lipofuscin accumulates in postmitotic cells with age, impairing their functioning. Unbalanced cell metabolic and waste-degradation functions cause its presence. IO neurons are relatively immune to apoptosis (Lasn et al., 2001); they preserve their function with aging, although they tend to accumulate significant amount of lipofuscin with age (Brizzee et al., 1975). It is unclear whether the presence of a large amount of lipofuscin is a result of higher lipofuscin generation and/or decelerated removal (Fonseca et al., 2005). Because lipofuscin aggregates are unavoidable reactions in biological systems, the lack of a cycle involving lipofuscin elimination is more plausible than the absence of lipofuscin generation (Yin, 1996). The r-aminobutyric acid scavenging effects proposed by Deng et al. (2010) over advanced lipoxidation end products (ALEs) may be instrumental in lipofuscin clearance in the olivary system. A gradual decline of r– aminobutyric acid presence with age may explain the accumulation of lipoxidation products in IO neurons. MVN GABAergic afferents are the main source of r–aminobutyric acid for the olivary cells, but they also mediate the electrical coupling among them. The gradual degeneration of these GABAergic afferents may explain the gradual presence of IO lipofuscin as well as the altered activations of IO ensembles with aging.

## Methods

### Vestibulo-Ocular Reflex (VOR) Model

The VOR was defined as a continuous-time mathematical model with two poles (Eq. 1), whose parameters were adjusted recursively to fit experimental and clinical data (Gordon et al., 1989; Gandhi et al., 2000):

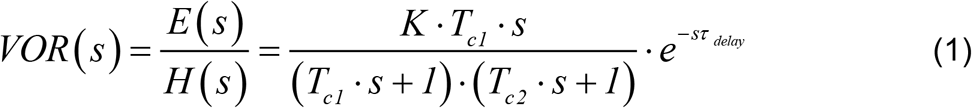

Where *e*(*t*), *E* (*s*) : *eye motion* (*output*), and *h*(*t*), *H* (*s*) : *head motion* (*input*)

There were 4 parameters in the model: *Q* = [*K*, T_*c*1_, T_*c*2_, *τ*_*delay*_]. The delay parameter *τ* _*delay*_ captured the delay in communicating the signals from the inner ear to the brain and the eyes. This delay is the consequence of the time needed for neurotransmitters to traverse the synaptic clefts between nerve cells. Based on the number of synapses involved in the VOR, the estimate of this delay is of 5 ms (Skavenski & Robinson, 1973; Robinson, 1981). The gain parameter *K*, assumed to be between 0.6 and 1, modelled the fact that the eyes do not perfectly cope with the movement of the head (Skavenski & Robinson, 1973; Robinson, 1981). The *T*_*c*1_ parameter represented the dynamics associated with the semicircular canals as well as some additional neural processing. The canals are high-pass filters, as the neural active membranes in the canals slowly relax back to their resting position after rotational experimentation (the canals stop sensing motion). Based on the mechanical characteristics of the canals, combined with additional neural processing which prolongs this time constant to improve the VOR accuracy, the *T*_*c*1_ parameter was estimated to be around 15 sec, in agreement with the biologically range which is 10-30 sec (Skavenski & Robinson, 1973; Robinson, 1981). Finally, the *T*_*c* 2_ parameter captured the dynamics of the oculomotor plant, i.e. the eye and the muscles and tissues attached to it. Its value was between 0.005 and 0.05 sec.

To find the temporal response for the VOR transfer function, we needed to calculate the inverse Laplace transform (Eq. 2). The outcome of the inverse Laplace transform consisted in a differential equation system defined in the same time domain as the spiking cerebellar network (see below; note that we modelled the delay and we inserted within the sensorimotor delay).

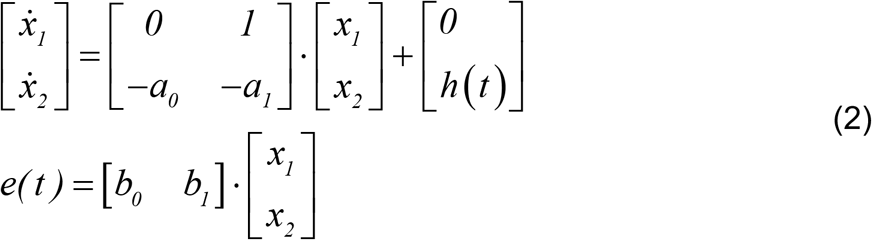

Where:

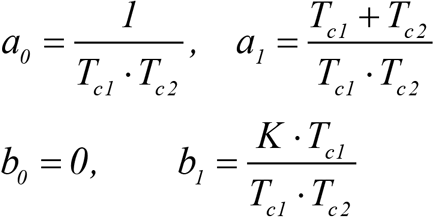

#### VOR analysis and assessment

The periodic functions representing eye and head velocities were analyzed through a discrete-time Fourier transform:

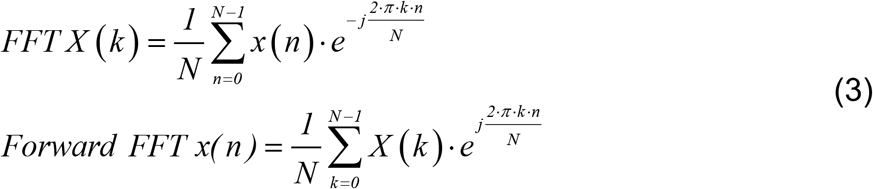

where *x* (*n*) indicates the periodic function, and *N* the number of samples within the considered time window. For each *k*, the term constituted a harmonic component (the complex version) with amplitude and frequency defined as:

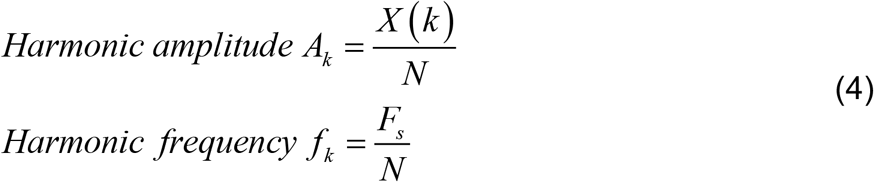

with *F*_*s*_ denoting the sampling frequency (0.5 KHz). The harmonic distortion values, which indicated the harmonic content of a waveform compared to its fundamental, were negligible. We calculated the **VOR gain** as the ratio between the first harmonic amplitudes of the forward Fourier eye- and head-velocity transforms

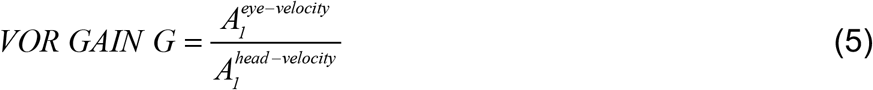

#### VOR protocols

In rotational chair testing, the subject (mouse, monkey, human) is seated on a rotatory table (Dumas et al., 2016). Speed and velocity of rotation are controlled and measured. The subject’s head is restrained, assuming that the movement of the table equals to the subject’s head movement. During normal VOR adaptation, a visual target is given in anti-phase with vestibular stimulation. The eyes must follow the visual target thus minimizing the retinal slip. In the model, the eye output function was defined as:

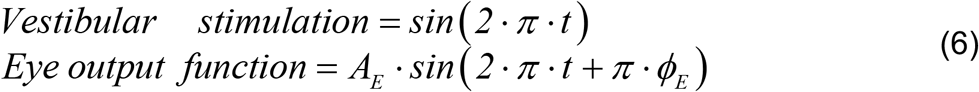

where the ideal VOR experiment values corresponded to *A*_*E*_ = −1,*ϕ*_*E*_ = 0 (visual field fixed).

### Cerebellar Network Model

The cerebellar network model consisted of five neural populations (Fig. 1C).

#### Mossy fibers (MFs)

100 MFs constituted the input to the cerebellar network. Mossy fibers (MFs) conveyed the sensory signals from the vestibular organ and the eye muscles onto granule cells (GCs) and medial vestibular nuclei (MVN). MF activity evolved based on a sinusoidal function (1Hz with a step size of 0.002 ms) to encode head movements consistently with the functional principles of VOR control (Lisberger & Fuchs, 1978; Arenz et al., 2008; Clopath et al., 2014; Badura et al., 2016). MF responses consisted of non-overlapping activations of equally sized neural subpopulations, which maintained a constant overall firing rate (Luque et al., 2016).

#### Granular cells (GCs)

2000 GCs operated as a state generator (Yamazaki & Tanaka, 2005, 2007, 2009). In the presence of a constant MF input, the granular layer generated a sequence of non-overlapping spatiotemporal patterns (i.e., states; Fujita, 1982). The same sequence of 500 states (each consisting of 4 active GCs per time step of 2 ms) repeatedly activated every 1-sec during learning (see below).

#### Purkinje cells (PCs)

We modelled a population of 200 PCs, divided into 2 groups of 100 cells to control agonist and antagonist eye muscles, respectively. PCs integrated the excitatory input from the parallel fibers (PFs), i.e. the axons of GCs, as well as the input from the climbing fibers (CFs), i.e. the axons of inferior olive (IO) cells. PCs projected inhibitory connections onto MVN cells, to close the cerebellar loop and generate the VOR output.

#### Inferior olive (IO) and climbing fibers (CFs)

We modelled 200 IO cells, divided in 2 groups of 100 IO cells for agonist/antagonist muscles, respectively. Each IO cell projected a CF onto one PC and one MVN cell. IO cells were interconnected via excitatory gap junctions, whose electrical coupling followed preferred directions (Devor & Yarom, 2002). The preferred paths were disposed radially from the center of 5×5 IO cell subpopulations, as in a square regular lattice network (Nobukawa & Nishimura, 2016). The strength of the electrical coupling, which drove the recurrent dynamics of the olivary population, was equal between all IO cells of the lattice network (see Table 1). In terms of external inputs, the IO population received excitatory afferents coding for retina slips (Clopath et al., 2014).This input reached the center of each lattice network and it was generated by a Poisson spiking process (Boucheny et al., 2005; Luque et al., 2011b). The IO population also received an inhibitory external input from MVN cells (Fig. 1C) whose action regulated the IO network synchronization via electrical coupling modulation (Best & Regehr, 2009; Lefler et al., 2014). We assumed a progressive age-related decrease of this inhibitory action based on the progressive age-loss of MVN neurons (Torvik et al., 1986), which modulated the MVN-IO inhibitory synaptic weight distribution of each 5×5 IO cell subpopulation. The variance of the Gaussian MVN-IO weight distribution varied linearly from 0.4 to 1.75 causing a more homogeneous electrical coupling along each 5×5 IO cell subpopulation whilst aging.

**Table 1.** Cerebellar network topology parameters.

| **Neurons** |  | **Synapses** |  |  |  |
| --- | --- | --- | --- | --- | --- |
| **Pre-synaptic cells (number)** | **Post-synaptic cells (number)** | **Number** | **Type** | **Initial weight** | **Weight range** |
| **2000 GCs** | **200 PCs** | **400000** | **AMPA** | **rand** | **[0, 3.65]** |
| **200 IO** | **200 PCs** | **200** | **AMPA** | **40** | **–** |
| **100 MFs** | **200 MVN** | **20000** | **AMPA** | **0** | **[0, 1]** |
| **200 PCs** | **200 MVN** | **200** | **GABA** | **1.5** | **–** |
| **200 IO** | **200 MVN** | **200** | **NMDA** | **7** | **–** |
| **IO to IO (lattice configuration)** |  | **320** | **EC** | **5** | **–** |

The error-related inputs (coding for retina slips), combined with the recurrent electrical coupling modulated by inhibitory MVN inputs, determined the overall activity of the IO population, which generated the CF bursting output. The probabilistic spike sampling of retina slips ensured an exploration of the whole error space over trials, whilst maintaining the CF activity below 10 Hz per fiber (in agreement with electrophysiological data; Kuroda et al., 2001). The evolution of the error could be sampled accurately even at such a low frequency (Carrillo et al., 2008; Luque et al., 2011b). A graded representation of the error signal (Najafi & Medina, 2013) led to a correlation between the intensity of the sampled instantaneous error and the number of the spikes within the CF burst (Eq. 7):

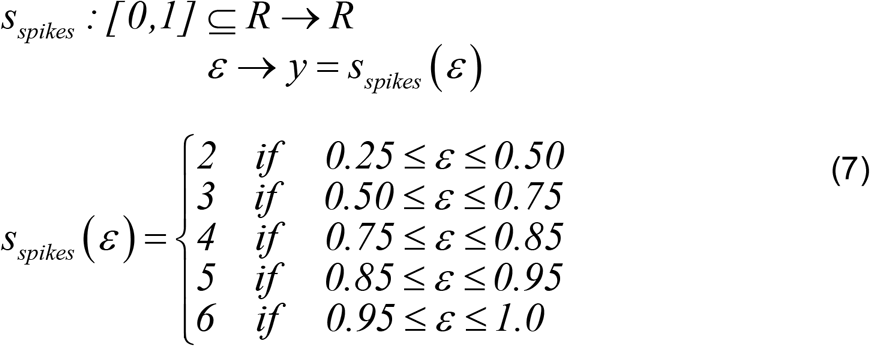

We assumed a perfect transmission of bursts from CFs to target PCs, i.e. the number of spikes in a PC complex spike linearly depended on the number of spikes in the CF burst (Mathy et al., 2009). The IO transmitted from 2 to 6 CF stimuli, delivered at inter-stimulus intervals of 2 ms, a range representative of inter-spike intervals recorded in olivary axons during bursts (Davie et al., 2008; Mathy et al., 2009), depending on the retina slips to be compensated.

#### Medial Vestibular Nuclei (MVN) cells

We modelled a population of 200 MVN cells, with again 2 groups of 100 cells for agonist/antagonist muscles, respectively. Each MVN cell received an inhibitory afferent from a PC and an excitatory afferent from the IO cell that was also contacting that PC (Uusisaari & De Schutter, 2011; Luque et al., 2014). MVN cells also received excitatory projections from all MFs. The subcircuit IO-PC-MVN was then organized in a single microcomplex. This circuitry arrangement rested upon the principles of circuit integrity and uniformity on the olivo-cortico-nucleo-olivary loop (Uusisaari & De Schutter, 2011).

#### Translation of MVN spike trains into analogue eye motor commands

The MVN output was translated into analogue output signals by averaging the spiking activity of each MVN subpopulation (one for each agonist/antagonist group of muscles) (Eqs. 8, 9):

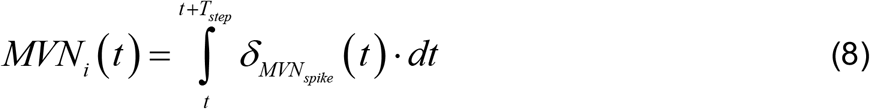

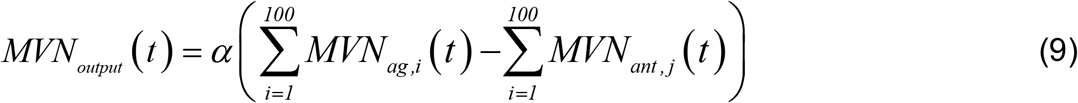

where *α* is the kernel amplitude that normalized the contribution of each MVN cell spike to the cerebellar output correction (the *MVN*_*ag*_ output controlled the agonist muscle, whilst the *MVN*_*ant*_ output controlled the antagonist muscle).

Table 1 summarizes the parameters of the cerebellar topology used in the model.

### Neuronal Models

#### MVN cell model

We modelled MVN cells as LIF neurons with excitatory (AMPA and NMDA) and inhibitory (GABA) chemical synapses (Eqs. 10-16).

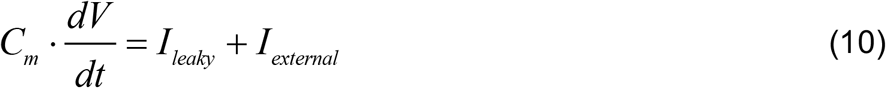

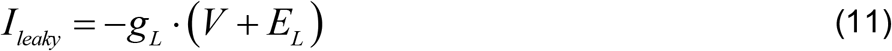

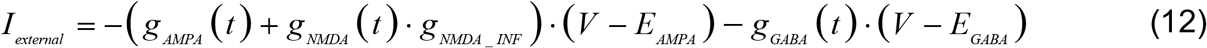

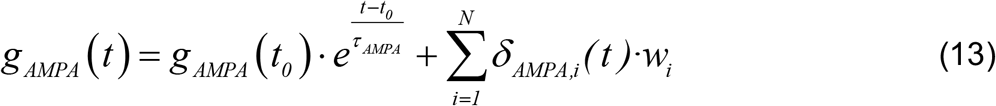

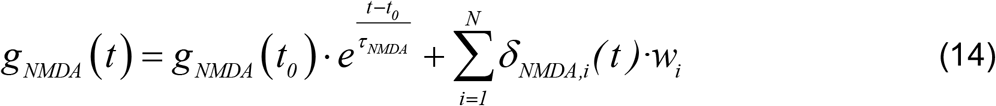

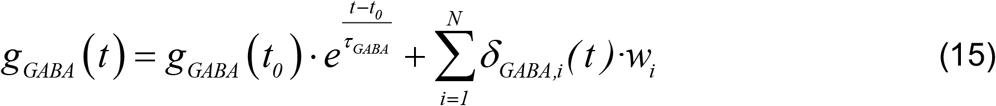

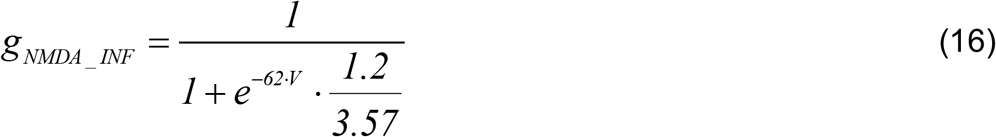

where: *C*_*m*_ denoted de membrane capacitance; *V* the membrane potential; *I*_*leaky*_ the leak current; *I*_*external*_ the external currents; *E*_*L*_ the resting potential; *g*_*L*_ the conductance responsible for the passive decay term towards the resting potential; *w*_*i*_ the synaptic weight of the synapses between the neuron *i* and the target neuron. Conductances *g*_*AMPA*_, g_*NMDA*_, *and g*_*GABA*_ integrated all the contributions received by each receptor (AMPA, NMDA, and GABA) through individual synapses. These conductances were defined as decaying exponential functions, which were proportionally incremented via *w*_*i*_ upon each presynaptic spike arrival (Dirac delta function). Finally, *g*_*NMDA* _INF_ stand for the NMDA activation channel. Note that we set the neuron membrane potential to *E*_*L*_ during the refractory period *(T*_*ref*_ ), just after reaching *V*_*thr*_ (voltage firing threshold) (Gerstner & Kistler, 2002; Gerstner et al., 2014). All the parameters of the neuronal models are shown in Table 2.

**Table 2.**
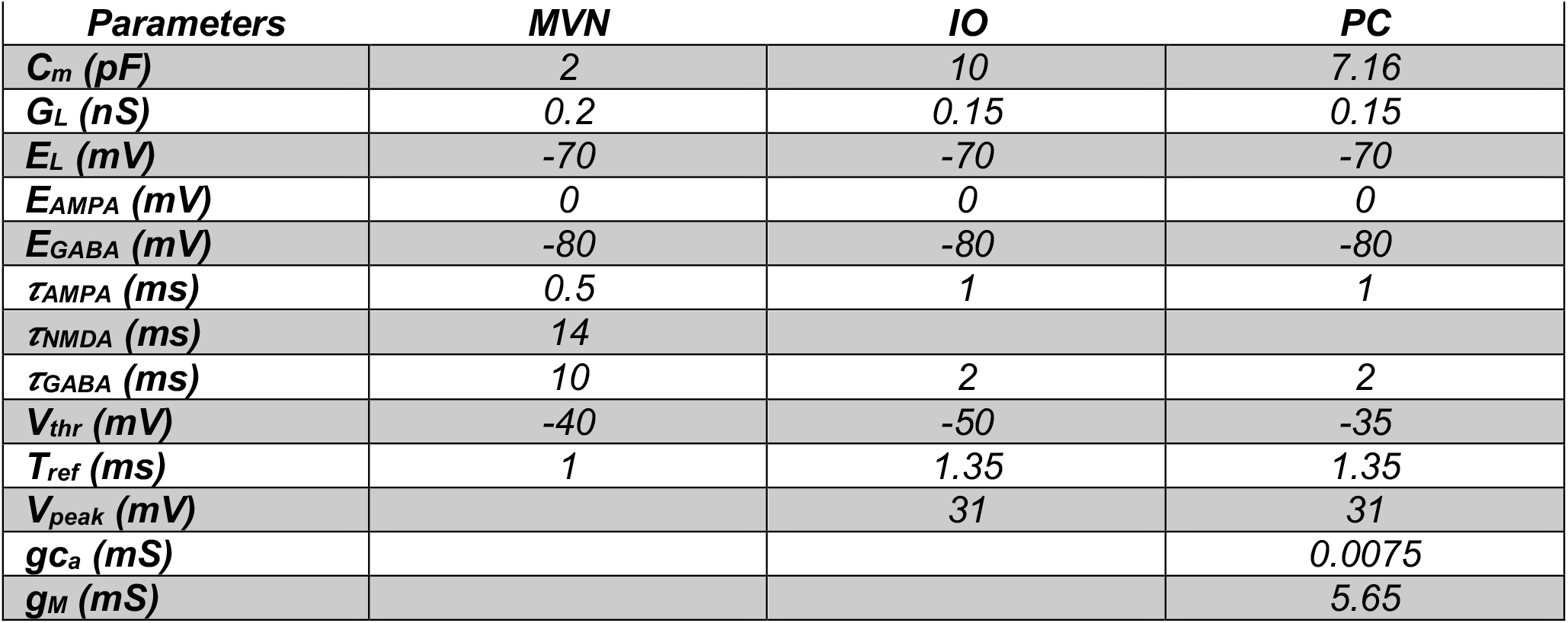
Neuronal model parameters.

| **Parameters** | **MVN** | **IO** | **PC** |
| --- | --- | --- | --- |
| **$C_m$ (pF)** | **2** | **10** | **7.16** |
| **$G_L$ (nS)** | **0.2** | **0.15** | **0.15** |
| **$E_L$ (mV)** | **-70** | **-70** | **-70** |
| **$E_{AMPA}$ (mV)** | **0** | **0** | **0** |
| **$E_{GABA}$ (mV)** | **-80** | **-80** | **-80** |
| **$\tau_{AMPA}$ (ms)** | **0.5** | **1** | **1** |
| **$\tau_{NMDA}$ (ms)** | **14** |  |  |
| **$\tau_{GABA}$ (ms)** | **10** | **2** | **2** |
| **$V_{thr}$ (mV)** | **-40** | **-50** | **-35** |
| **$T_{ref}$ (ms)** | **1** | **1.35** | **1.35** |
| **$V_{peak}$ (mV)** |  | **31** | **31** |
| **$g_{Ca}$ (mS)** |  |  | **0.0075** |
| **$g_M$ (mS)** |  |  | **5.65** |

#### Inferior olive (IO) neuronal model

We modelled IO cells as LIF neurons with excitatory (AMPA) and inhibitory (GABA) chemical synapses as well as with electronic gap junctions (Llinas et al., 1974; Sotelo et al., 1974) (Eqs. 17-22):

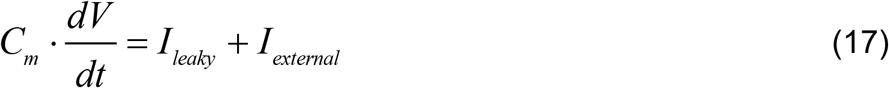

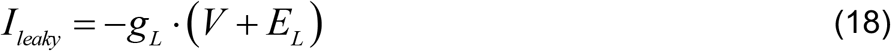

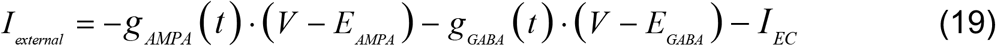

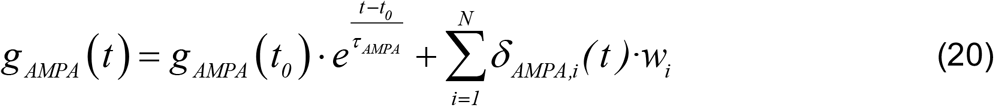

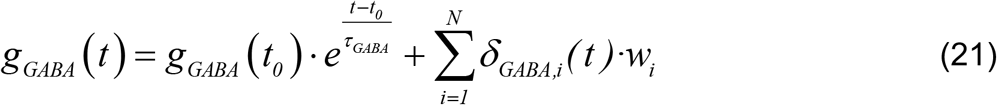

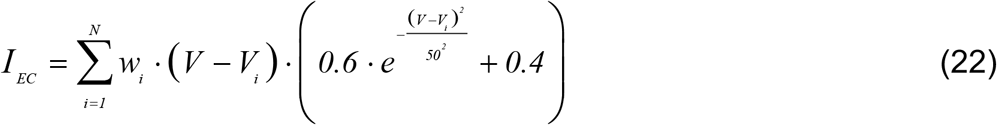

where: *C*_*m*_ denotes de membrane capacitance; *V* the membrane potential; *I*_*leaky*_ the leak current; *I*_*external*_ *l* the external currents; *E*_*L*_ the resting potential; *g*_*L*_ the conductance responsible for the passive decay term toward the resting potential; *w*_*i*_ the synaptic weight of the synapses between the neuron *i* and the target neuron. Conductances *g*_*AMPA*_ *and g*_*GABA*_ integrated all the contributions received by each chemical receptor (AMPA, GABA) through individual synapses. These conductances were defined as decaying exponential functions, which were proportionally incremented via *w*_*i*_ upon each presynaptic spike arrival (Dirac delta function) (Gerstner & Kistler, 2002; Ros et al., 2006). *I*_*EC*_ represented the total current injected through the electrical synapses (Schweighofer et al., 1999). *V* was the membrane potential of the target neuron, *V*_*i*_ the membrane potential of the neuron *i, and N* was the total number of input synapses of the target neuron. Finally, for a correct operation of the electrical synapses, this model emulated the depolarization and hyperpolarization phases of an action potential. The LIF neuron incorporated a simple threshold process that enabled the generation of a triangular voltage function (maximum/minimum value *V*_*peak*_ *E*_*L*_ respectively) each time the neuron fired (Bezzi et al., 2004). All the parameters of the IO neuronal model are shown in Table 2.

#### Purkinje cell model

The PC model was the same as in (Miyasho et al., 2001; Middleton et al., 2008; Luque et al., 2019). It reproduced the three spiking modes of Purkinje cells, namely tonic, bursting, and spike pauses (Forrest, 2008). The PC model consisted of a single compartment with five ionic currents and two excitatory (AMPA) and inhibitory (GABA) chemical synapses (Eqs. 23-27):

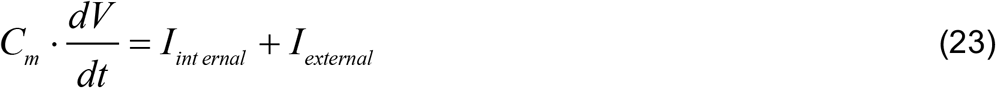

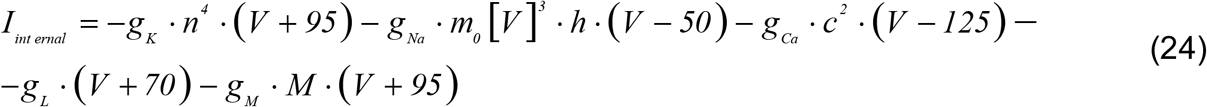

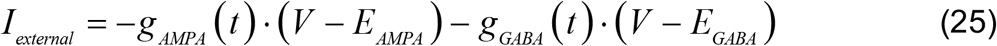

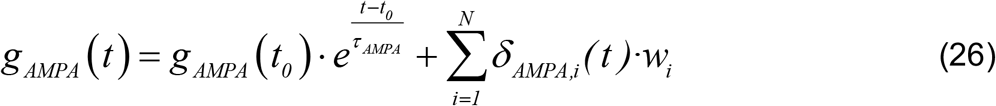

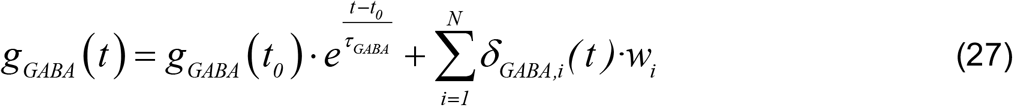

where *C*_*m*_ denotes de membrane capacitance, *V* the membrane potential, *I* the internal currents, *I*_*external*_ the external currents, and *w*_*i*_ the synaptic weight of the synapses between the neuron *i* and the target neuron. Conductances *g*_*AMPA*_ *and g*_*GABA*_ integrated all the contributions received by each chemical receptor type (AMPA, GABA) through individual synapses. These conductances were decaying exponential functions that were proportionally incremented via *w*_*i*_ upon each presynaptic spike arrival (Dirac delta function) (Gerstner & Kistler, 2002; Ros et al., 2006). Finally, *g*_*K*_ was the delayed rectifier potassium current, *g*_*Na*_ the transient inactivating sodium current, *g*_*Ca*_ the high-threshold non-inactivating calcium current, *g*_*L*_ the leak current, and *g*_*M*_ the muscarinic receptor suppressed potassium current (see Table 3).

**Table 3.**
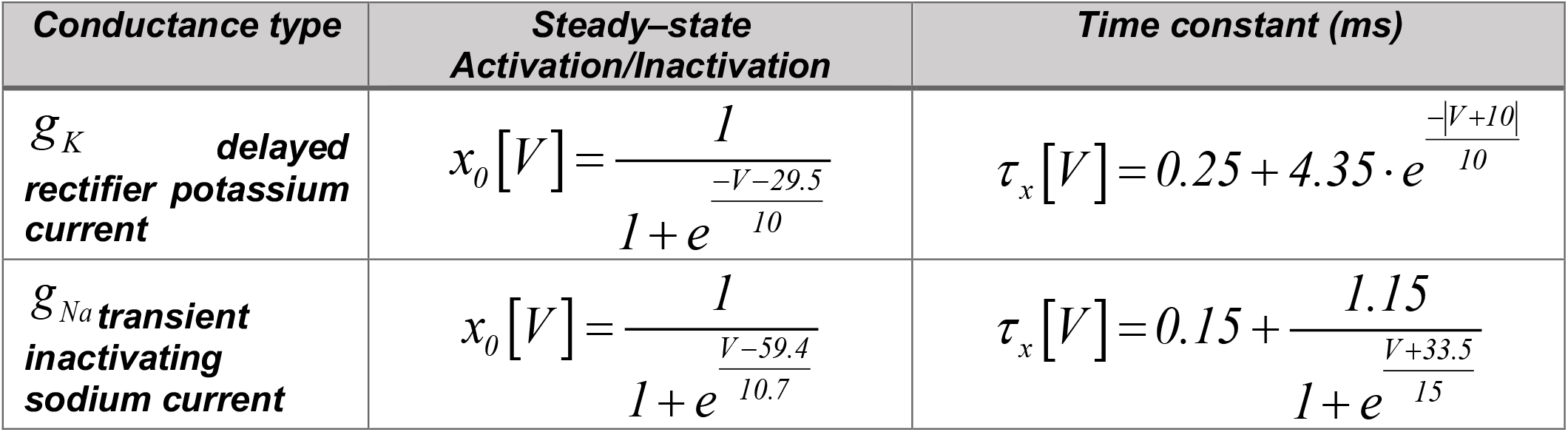
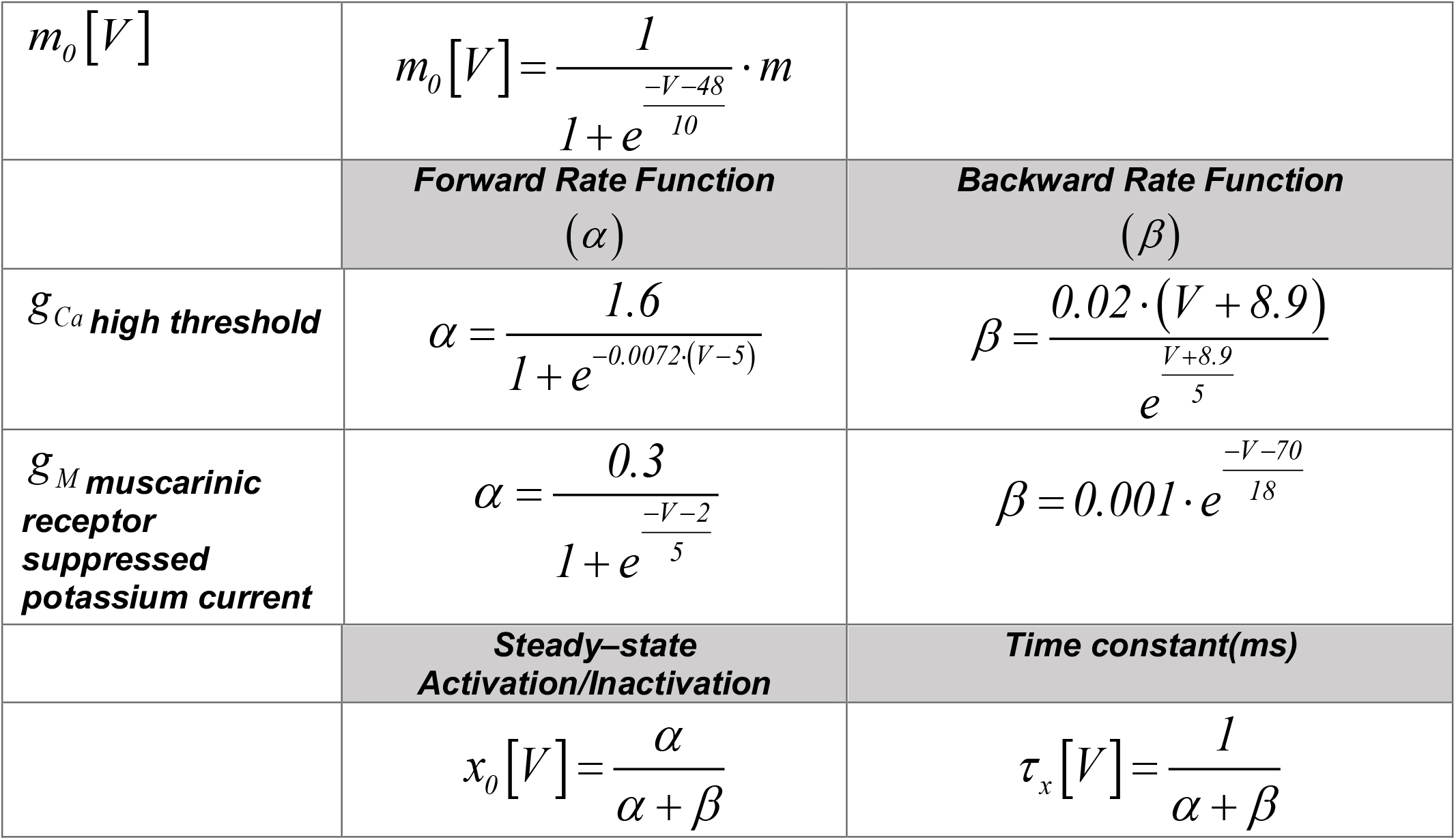
Ionic conductance kinetic parameters.

The dynamics of each gating variable (*n, h, c*, and *M*) followed the Eq. 28:

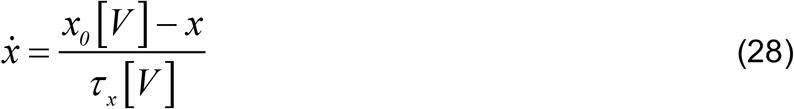

where *x* corresponds to variables *n, h, c*, and *M*. The equilibrium function was given by the term *x*_*0*_ [*V*] and the time constant *τ* _*x*_ [*V*] (see Table 3).

The sodium activation variable was replaced and approximated by its equilibrium function *m*_*0*_ [*V*]. The *M* current presented a temporal evolution significantly slower than the rest of variables. Each spike in the neuron generated a fast increase of the *M* current that took several milliseconds to return to its stable state. A high *M* current prevented the PC from entering in its tonic mode (when the neuron generated spikes due to PFs activity). A complex spike caused a rapid increase of the *M* current that depended, in turn, on the size of the spikelet within the burst. PC tonic mode resumed when the *M* current decreased.

We first validated the PC model in the NEURON simulator and then we reduced it to make it compatible with EDLUT (Luque et al., 2019). In the reduced PC model, we implemented the *I*_*K*_ and *I*_*na*_ currents through a simple threshold process that triggered the generation of a triangular voltage function each time the neuron fired (Bezzi et al., 2004). This triangular voltage depolarization drove the state of ion channels similarly to the original voltage depolarization during the spike generation. The final internal current was given by Eq. 29:

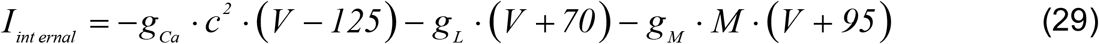

All the parameters are shown in Table 3.

#### Mossy fibers (MF) & granule cells (GC) models

MFs and GC neurons were simulated as leaky integrate–and–fire (LIF) neurons, with the same excitatory (AMPA) and inhibitory (GABA) chemical synapses and parameters as in Luque et al. (2019).

### Synaptic Plasticity Models

#### PC Intrinsic Plasticity

We equipped the PC model with a mechanism to update the value of the membrane capacitance (Cm) according to Eq. 30:

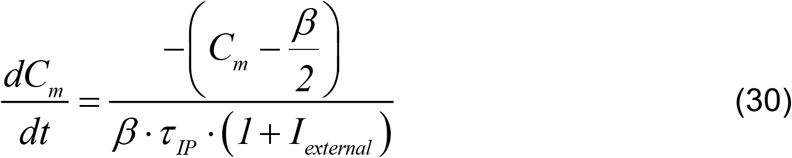

where *τ* _*IP*_ denotes the intrinsic plasticity time constant set to 12.10^3^ sec (this large time constant prevented interferences between intrinsic plasticity and other STDP mechanisms during the learning process (Garrido et al., 2016); *β* controls the shape of the firing rate distribution and it is equal to 1 (see Garrido et al., 2016) for details about all intrinsic plasticity mechanism parameters). Whenever a spike was elicited, the *C*_*m*_ variable was updated according to the following equation:

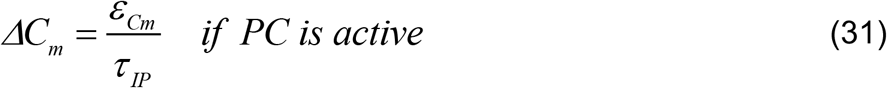

where *ε*_*Cm*_ = *0*.*0475* (Garrido et al., 2016) determined the influence of each spike on *C*_*m*_. Note that the membrane capacitance of PCs could not diminish below a lower limit (0.77 ± 0.17 μF cm^−2^ where mean ± s.d.; range, 0.64-1.00 μF cm^−2^; Roth & Häusser, 2001).

#### PF-PC synaptic plasticity

The model of long-term depression (LTD) and long-term potentiation (LTP) at PF–PC synapses was the same as in Luque et al. (2019), and it followed the Eqs. 32 and 33:

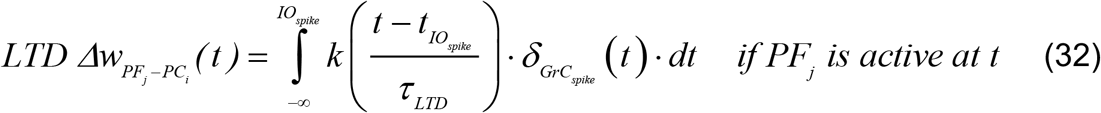

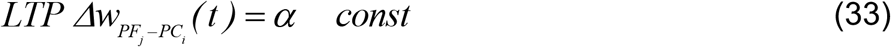

where 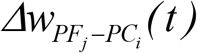 denotes the weight change between the *j*^*th*^ PF and the target *i^th^* PC;*τ* _*LTD*_ is the time constant that compensates for the sensory motor delay (i.e., about 100 ms; Sargolzaei et al., 2016); *δ*_*GrC*_ is the Dirac delta function corresponding to an afferent spike from a PF; and the kernel function *k* (*x*) is defined as in Eq. 34:

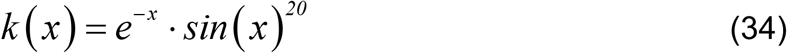

With this parametric configuration, the effect on presynaptic spikes arriving through PFs is maximal over the 100 ms time window before CF spike arrival, thus accounting for the sensorimotor pathway delay. For the sake of computational efficiency, note that the kernel *k* (*x*)combines exponential and trigonometric functions that allow for recursive computation suitable for an event-driven simulation scheme as EDLUT (Ros et al., 2006; Naveros et al., 2015; Naveros et al., 2017). Computational recursion avoids integrating the whole kernel upon each new spike arrival.

Finally, as shown in Eq. 33, the amount of LTP at PFs was fixed (Kawato & Gomi, 1992; Luque et al., 2011a; Luque et al., 2016), with an increase of synaptic efficacy equal to α each time a spike arrived through a PF to the targeted PC. This STDP mechanism correlated the activity patterns coming through the PFs to PCs with the instructive signals coming from CFs to PCs (producing LTD in the activated PF–PC synapses). The correlation process at PC level identified certain PF activity patterns and it consequently reduced the PC output activity. A decrease of PC activations caused a subsequence reduction on the PC inhibitory action over the target MVN. Since the MVN received an almost constant gross MF activation, a lack of PC inhibitory action caused increasing levels of MVN activation. Conversely, the STDP mechanism increased the PC inhibitory activity by potentiating PF-PC synapses in the absence of instructive signal, thus causing decreasing levels of MVN activations. Consequently, PC axon activity governed MVN activation by shaping their inhibitory action produced onto MVN. This spike-timing-dependent plasticity (STDP) mechanism, which regulated the LTP/LTD ration at PF-PC synapses, shaped the inhibitory action of PCs onto MVN cells.

#### MF-MVN synaptic plasticity

The LTD/LTP dynamics at MF-MVN synapses were the same as in (Luque et al., 2019)), i.e., they were based on the following rules:

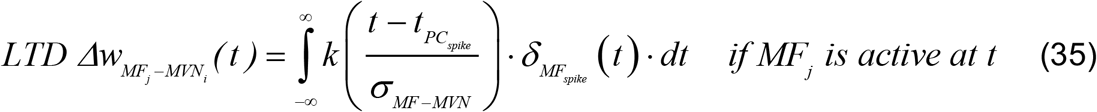

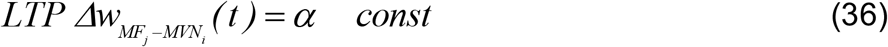

with 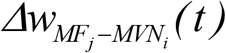 denotes the weight change between the *j*^*th*^ MF and the target *i^th^* MVN; *σ* _*MF* −*MVN*_ the temporal width of the kernel; and *δ*_*MF*_ the Dirac delta function that defined a MF spike. The integrative kernel function *k* (*x*) was taken as:

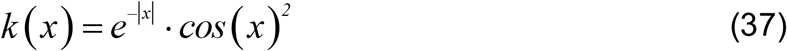

Note that there is no need for sensorimotor delay compensation thanks to the previous learning rule (*τ* _*LTD*_ in Eq. 32). This second STDP mechanism accounted for learning consolidation at MVN (see Luque et al., 2016). The PC output operated as an instructive signal and correlated the activity patterns coming from MFs to MVN (producing LTD in the activated MF–MVN synapses upon the arrival of the instructive signal and LTP otherwise). Well-timed sequences of increasing/decreasing levels of MVN activation ultimately shaped the cerebellar output during VOR adaptation.

The EDLUT source code is available at the following URL:

www.ugr.es/~nluque/restringido/CODE_Cerebellar_Ageing_Vestibulo_Ocular_Adaptation.rar

User: REVIEWER, password: REVIEWER.

## Figure Captions

**Supplementary Figure 1.**
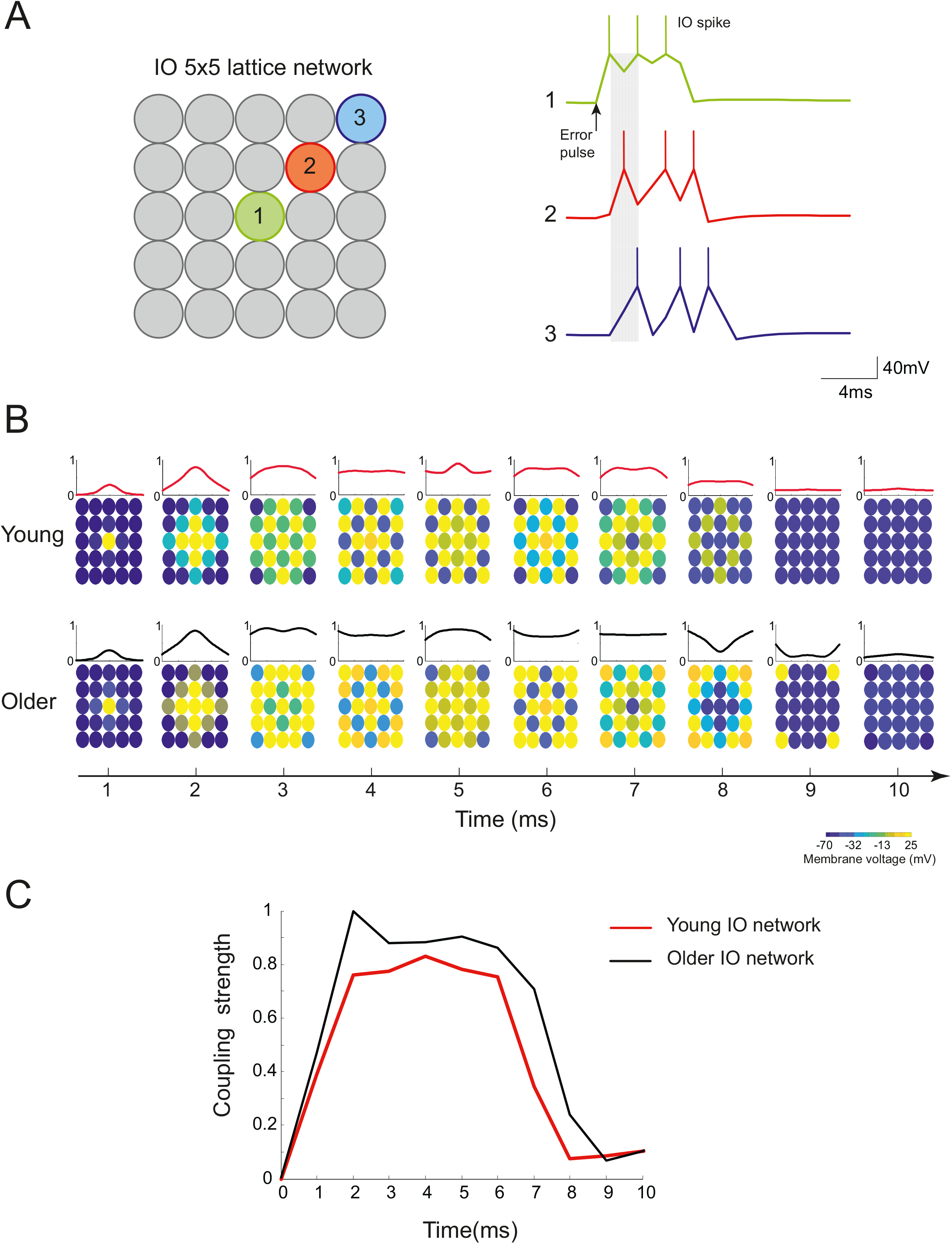
Spatial-temporal evolution of IO network activity patterns in young and older individuals. **(A)** Spike propagation mediated by electrical coupling along the diagonal of an IO cell cluster (5×5 lattice configuration). Spike activity was induced by an external stimulus delivered at the center of the network. **(B)** Time course of the membrane potentials of IO cells in 5×5 clusters induced by a large input stimulus received at the center of the network at 1 ms. Top: young individuals. Bottom: older individuals. **(C)** Temporal profile of the strength of the electrical coupling between IO cells in young (red curve) and older (black curve) individuals.

**Supplementary Figure 2.**
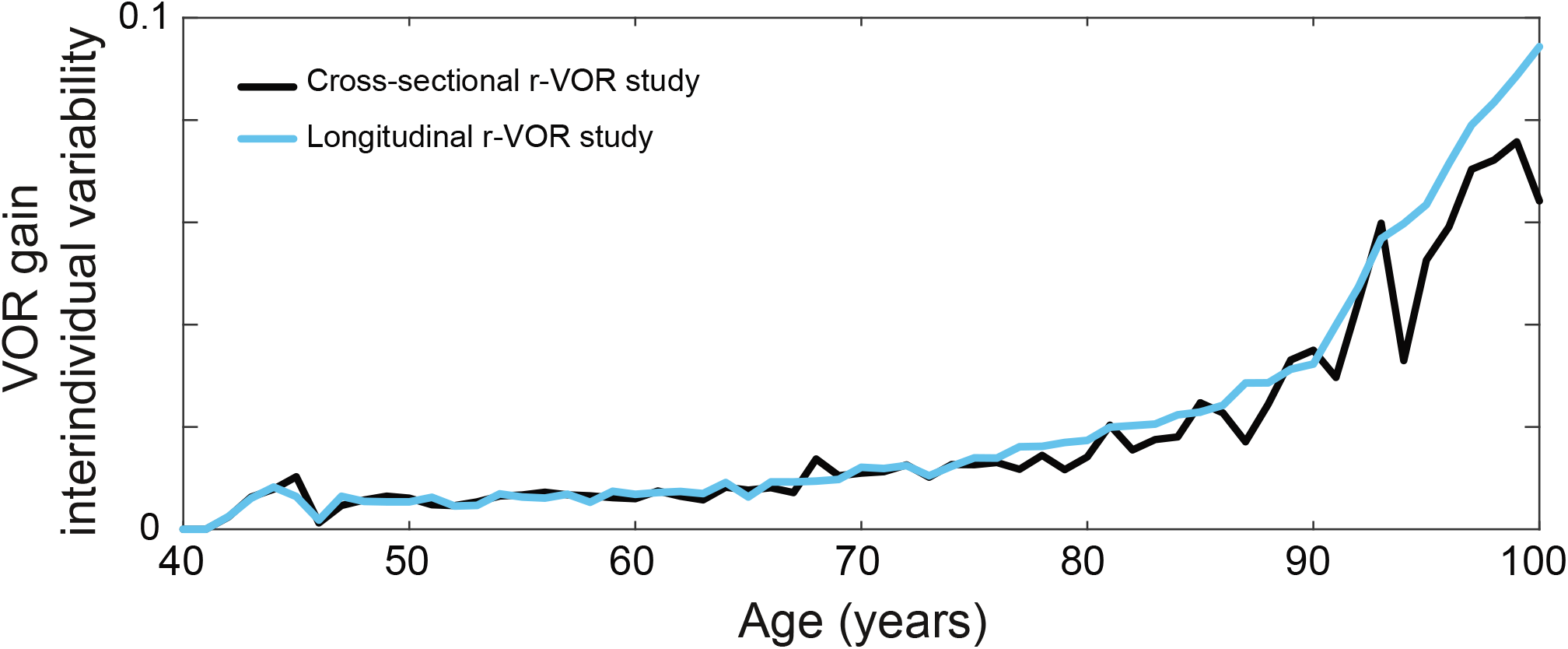
Interindividual variability of r-VOR gain as a function of age. Black curve: r-VOR standard deviation of the variance across individuals recorded during the cross-sectional aging simulation (Fig. 6A). Light-blue curve: r-VOR standard deviation of the variance across individuals recorded during the longitudinal aging simulation (Fig. 7A).

**Supplementary Figure 3.**
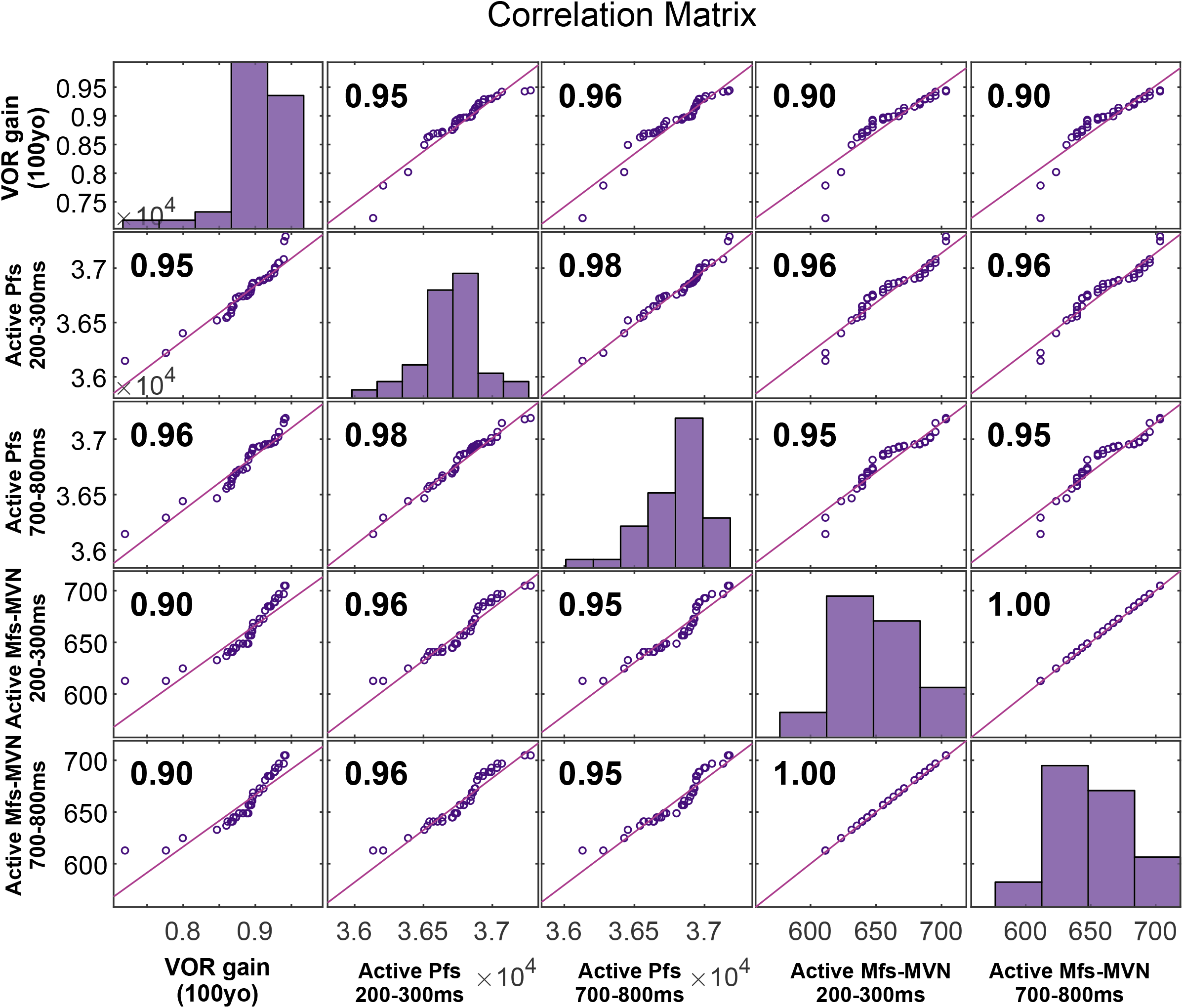
Correlation matrix between VOR gain and the number of residual PFs and MF-MVN projections active at the peak and the trough of the eye velocity function. The diagonal represents the histograms of the VOR performance values.

## References

Allen, D., Ribeiro, L., Arshad, Q., & Seemungal, B. M. (2017). Age-related vestibular loss: current understanding and future research directions. Front Neurol., 7, 231.

Alvarez, J. C., Diaz, C., Suarez, C., Fernandez, J. A., del Rey, C. G., Navarro, A., & Tolivia, J. (2000). Aging and the human vestibular nuclei: morphometric analysis. Mechanisms Ageing & Development, 114(3), 149–172.

Andersen, B. B., Gundersen, H. J. G., & Pakkenberg, B. (2003). Aging of the human cerebellum: a stereological study. J. Comparative Neurol., 466(3), 356–65.

Anson, E., & Jeka, J. (2016). Perspectives on aging vestibular function. Front. Neurol., 6, 269.

Arenz, A., Silver, R. A., Schaefer, A. T., & Margrie, T. W. (2008). The Contribution of Single Synapses to Sensory Representation in Vivo. Science, 321(5891), 977–980.

Badura, A., Clopath, C., Schonewille, M., & De Zeeuw, C. I. (2016). Modeled changes of cerebellar activity in mutant mice are predictive of their learning impairments. Sci. Rep., 6, 36131.

Baloh, R. W., Jacobson, K. M., & Socotch, T. M. (1993). The effect of aging on visual-vestibuloocular responses. Exp. Brain Res., 95(3), 509–516.

Baloh, R. W., Sloane, P. D., & Honrubia, V. (1989). Quantitative vestibular function testing in elderly patients with dizziness. Ear, nose, & throat journal, 68(12), 935–9.

Bergström, B. (1973). Morphology of the vestibular nerve: III. Analysis of the calibers of the myelinated vestibular nerve fibers in man at various ages. Acta oto-laryngol., 76(1-6), 331–38.

Best, A. R., & Regehr, W. G. (2009). Inhibitory regulation of electrically coupled neurons in the inferior olive is mediated by asynchronous release of GABA. Neuron, 62(4), 555–65.

Bezzi, M., Nieus, T., Coenen, O. J.-M., & D’Angelo, E. (2004). An I&Fmodel of a cerebellar granule cell. Neurocomputing, 58, 593–8.

Boucheny, C., Carrillo, R. R., Ros, E., & Coenen, O. J.-M. D. (2005). Real-time spiking neural network: an adaptive cerebellar model. LNCS, 3512, 136–144.

Brandt, T., Schautzer, F., Hamilton, D. A., Brüning, R., Markowitsch, H. J., Kalla, R., Darlington, C., Smith, P., & Strupp, M. (2005). Vestibular loss causes hippocampal atrophy and impaired spatial memory in humans. Brain, 128(11), 2732–41.

Brizzee, K. R., Kaack, B., & Klara, P. (1975). Lipofuscin: intra-and extraneuronal accumulation and regional distribution. In Neurobiol. of Aging (pp. 463–484). Springer.

Carrillo, R. R., Ros, E., Boucheny, C., & Coenen, O. J.-M. (2008). A real-time spiking cerebellum model for learning robot control. Biosystems, 94(1-2), 18–27.

Clopath, C., Badura, A., De Zeeuw, C. I., & Brunel, N. (2014). A cerebellar learning model of VOR adaptation in wild-type and mutant mice. J. Neurosci., 34(21), 7203–7215.

D’Angelo, E., & De Zeeuw, C. I. (2009). Timing and plasticity in the cerebellum: focus on the granular layer. Trends Neurosci., 32(1), 10.

D’Angelo, E., Mapelli, L., Casellato, C., Garrido, J. A., Luque, N.R., Monaco, J., Prestori, F., Pedrocchi, A., & Ros, E. (2016). Distributed circuit plasticity: new clues for the cerebellar mechanisms of learning. Cerebellum, 15(2), 139–151.

Davie, J. T., Clark, B. A., & Häusser, M. (2008). The origin of the complex spike in cerebellar Purkinje cells. J. Neurosci., 28(30), 7599–7609.

De Zeeuw, C. I., Hoogenraad, C. C., Koekkoek, S., Ruigrok, T. J., Galjart, N., & Simpson, J. I. (1998). Microcircuitry and function of the inferior olive. Trends Neurosci., 21(9), 391–400.

Demer, J. L., Honrubia, V., & Baloh, R. W. (1994). Dynamic visual acuity: a test for oscillopsia and vestibulo-ocular reflex function. The American Journal of Otology, 15(3), 340–7.

Deng, Y., Xu, L., Zeng, X., Li, Z., Qin, B., & He, N. (2010). New perspective of GABA as an inhibitor of formation of advanced lipoxidation end-products: it’s interaction with malondiadehyde. J. Biomed. Nanotechnol., 6(4), 318–24.

Desai, A., Goodman, V., Kapadia, N., Shay, B. L., & Szturm, T. (2010). Relationship between dynamic balance measures and functional performance in community-dwelling elderly people. Physical Therapy, 90(5), 748–60.

Devor, A., & Yarom, Y. (2002). Generation and propagation of subthreshold waves in a network of inferior olivary neurons. J Neurophysiol., 87(6), 3059–69.

Dits, J., Houben, M. M., & van der Steen, J. (2013). Three dimensional Vestibular ocular reflex testing using a Six degrees of freedom motion platform. Journal of visualized experiments: JoVE(75).

Dumas, G., Perrin, P., Ouedraogo, E., & Schmerber, S. (2016). How to perform the skull vibration-induced nystagmus test (SVINT). Eur Annals of Otorhinolaryngol., Head & Neck Diseases, 133(5), 343–8.

Fonseca, D., Sheehy, M., Blackman, N., Shelton, P., & Prior, A. (2005). Reversal of a hallmark of brain ageing: lipofuscin accumulation. Neurobiol. of Aging, 26(1), 69–76.

Forrest, M. (2008). Biophysics of Purkinje computation University of Warwick].

Fujita, M. (1982). Adaptive filter model of the cerebellum. Biological Cybernetics, 45(3), 195–206.

Gandhi, C. C., Kelly, R. M., Wiley, R. G., & Walsh, T. J. (2000). Impaired acquisition of a Morris water maze task following selective destruction of cerebellar purkinje cells with OX7-saporin. Behavioural Brain Res., 109(1), 37–47.

Gao, Z., vanBeugen, B. J., & De Zeeuw, C. I. (2012). Distributed Synergistic Plasticity and Cerebellar Learning. Nat. Rev. Neurosci., 13, 1–17.

Garrido, J. A., Luque, N. R., Tolu, S., & D’Angelo, E. (2016). Oscillation-driven spike-timing dependent plasticity allows multiple overlapping pattern recognition in inhibitory interneuron networks. Int. J. Neural Syst., 26(05).

Gerstner, W., & Kistler, W. M. (2002). Spiking neuron models: Single neurons, populations, plasticity. Cambridge university press.

Gerstner, W., Kistler, W. M., Naud, R., & Paninski, L. (2014). Neuronal dynamics: From single neurons to networks and models of cognition. Cambridge University Press.

Gordon, J., Furman, J., & Kamen, E. (1989). System identification of the vestibulo-ocular reflex: application of the recursive least-squares algorithm. Bioengineering Conference, 1989., Proceedings of,

Grasselli, G., He, Q., Wan, V., Adelman, J. P., Ohtsuki, G., & Hansel, C. (2016). Activity-Dependent Plasticity of Spike Pauses in Cerebellar Purkinje Cells. Cell Reports, 14(11), 2546–2553.

Grossman, G. E., & Leigh, R. J. (1990). Instability of gaze during locomotion in patients with deficient vestibular function. Annals of Neurology: Official Journal of the American Neurological Association and the Child Neurology Society, 27(5), 528–32.

Ichikawa, R., Sakimura, K., & Watanabe, M. (2016). GluD2 endows parallel fiber–Purkinje cell synapses with a high regenerative capacity. J. Neurosci., 36(17), 4846–58.

Ishikawa, T., Shimuta, M., & Häusser, M. (2015). Multimodal sensory integration in single cerebellar granule cells in vivo. Elife, 4, e12916.

Ito, M. (2013). Error Detection and Representation in the Olivo-Cerebellar System. Front. Neural Circuits, 1–8.

Jahn, K., Naeßl, A., Schneider, E., Strupp, M., Brandt, T., & Dieterich, M. (2003). Inverse U-shaped curve for age dependency of torsional eye movement responses to galvanic vestibular stimulation. Brain, 126(7), 1579–89.

Jang, D. C., Shim, H. G., & Kim, S. J. (2020). Intrinsic plasticity of cerebellar purkinje cells contributes to motor memory consolidation. J. Neurosci., 40(21), 4145–57.

Kawato, M., & Gomi, H. (1992). A computational model of four regions of the cerebellum based on feedback-error learning. Biol. Cybern., 68(2), 95–103.

Lasn, H., Winblad, B., & Bogdanovic, N. (2001). The number of neurons in the inferior olivary nucleus in Alzheimer’s disease and normal aging: A stereological study using the optical fractionator. J. Alzheimer’s Disease, 3(2), 159–68.

Latorre, R., Aguirre, C., Rabinovich, M. I., & Varona, P. (2013). Transient dynamics and rhythm coordination of inferior olive spatio-temporal patterns. Front. Neural Circuits, 7 138

Lefler, Y., Amsalem, O., Vrieler, N., Segev, I., & Yarom, Y. (2020). Using subthreshold events to characterize the functional architecture of the electrically coupled inferior olive network. Elife, 9, e43560.

Lefler, Y., Yarom, Y., & Uusisaari, M. Y. (2014). Cerebellar inhibitory input to the inferior olive decreases electrical coupling and blocks subthreshold oscillations. Neuron, 81(6), 1389–400.

Leigh, R. J., & Zee, D. S. (2015). The neurology of eye movements. Oxford University Press.

Li, C., Layman, A. J., Geary, R., Anson, E., Carey, J. P., Ferrucci, L., & Agrawal, Y. (2015). Epidemiology of vestibulo-ocular reflex function: data from the Baltimore Longitudinal Study of Aging. Otol. Neurotol., 36(2), 267–72.

Li, C., & Li, Y. (2013). A spike-based model of neuronal intrinsic plasticity. IEEE Trans. Autonomous Mental Development, 5(1), 62–73.

Lisberger, S. G., & Fuchs, A. F. (1978). Role of primate flocculus during rapid behavioral modification of VOR. II. Mossy fiber firing patterns during horizontal head rotation and eye movement. J. Neurophysiol., 41(3), 764–77.

Llinas, R., Baker, R., & Sotelo, C. (1974). Electrotonic coupling between neurons in cat inferior olive. J. Neurophysiol., 37(3), 560–71.

Loewenstein, Y. (2002). A possible role of olivary gap-junctions in the generation of physiological and pathological tremors. Molecular Psychiatry, 7(2), 129.

Lopez, I., Honrubia, V., & Baloh, R. W. (1996). Aging and the human vestibular nucleus. J. Vestibular Research: Equilibrium & Orientation, 7(1), 77–85.

Lorente de Nó, R. (1933). Vestibulo-ocular reflex arc. Archiv Neurol & Psychiatry.

Luque, N. R., Garrido, J. A., Carrillo, R. R., Coenen, O. J. M. D., & Ros, E. (2011a). Cerebellar Input Configuration Toward Object Model Abstraction in Manipulation Tasks. IEEE Trans. Neural. Netw., 22(8), 1321–28.

Luque, N. R., Garrido, J. A., Carrillo, R. R., Coenen, O. J. M. D., & Ros, E. (2011b). Cerebellarlike Corrective Model Inference Engine for Manipulation Tasks. IEEE Trans. Syst. Man. Cybern., 41(5), 1299–312.

Luque, N. R., Garrido, J. A., Carrillo, R. R., D’Angelo, E., & Ros, E. (2014). Fast convergence of learning requires plasticity between inferior olive and deep cerebellar nuclei in a manipulation task: a closed-loop robotic simulation. Front. Comput. Neurosci., 8.

Luque, N. R., Garrido, J. A., Naveros, F., Carrillo, R. R., D’Angelo, E., & Ros, E. (2016). Distributed Cerebellar Motor Learning; a STDP Model. Front. Comp. Neurosci., 10. https://doi.org/10.3389/fncom.2016.00017

Luque, N. R., Naveros, F., Carrillo, R. R., Ros, E., & Arleo, A. (2019). Spike burst-pause dynamics of Purkinje cells regulate sensorimotor adaptation. PLOS Comp.l Biol., 15(3), e1006298.

Mathy, A., Ho, S. S., Davie, J. T., Duguid, I. C., Clark, B. A., & Hausser, M. (2009). Encoding of oscillations by axonal bursts in inferior olive neurons. Neuron, 62(3), 388–99.

Matiño-Soler, E., Esteller-More, E., Martin-Sanchez, J. C., Martinez-Sanchez, J. M., & Perez-Fernandez, N. (2015). Normative data on angular vestibulo-ocular responses in the yaw axis measured using the video head impulse test. Otology & Neurotology, 36(3), 466–71.

McGarvie, L. A., MacDougall, H. G., Halmagyi, G. M., Burgess, A. M., Weber, K. P., & Curthoys, I. S. (2015). The video head impulse test (vHIT) of semicircular canal function–age-dependent normative values of VOR gain in healthy subjects. Front. Neurol., 6, 154.

Mergner, T., & Rosemeier, T. (1998). Interaction of vestibular, somatosensory and visual signals for postural control and motion perception under terrestrial and microgravity conditions—a conceptual model. Brain Res. Rev., 28(1-2), 118–35.

Middleton, S. J., Racca, C., Cunningham, M. O., Traub, R. D., Monyer, H., Knopfel, T., Schofield, I. S., Jenkins, A., & Whittington, M. A. (2008). High-frequency network oscillations in cerebellar cortex. Neuron, 58(5), 763–74.

Miyasho, T., Takagi, H., Suzuki, H., Watanabe, S., Inoue, M., Kudo, Y., & Miyakawa, H. (2001). Low-threshold potassium channels and a low-threshold calcium channel regulate Ca2+ spike firing in the dendrites of cerebellar Purkinje neurons: a modeling study. Brain Res., 891(1-2), 106–15.

Najac, M., & Raman, I. M. (2015). Integration of Purkinje Cell Inhibition by Cerebellar Nucleo-Olivary Neurons. J. Neurosci., 35(2), 544–9.

Najafi, F., & Medina, J. F. (2013). Beyond “all-or-nothing” climbing fibers: graded representation of teaching signals in Purkinje cells. Front. Neural Circuits, 7, 1–15.

Naveros, F., Garrido, J. A., Carrillo, R. R., Ros, E., & Luque, N. R. (2017). Event-and Time-Driven Techniques Using Parallel CPU-GPU Co-processing for Spiking Neural Networks. Front. Neuroinformatics, 11.

Naveros, F., Luque, N. R., Garrido, J. A., Carrillo, R. R., Anguita, M., & Ros, E. (2015). A Spiking Neural Simulator Integrating Event-Driven and Time-Driven Computation Schemes Using Parallel CPU-GPU Co-Processing: A Case Study. IEEE Trans. Neural Netw. Learn. Syst., 26(7), 1567–74.

Naveros, F., Luque, N. R., Ros, E., & Arleo, A. (2020). VOR Adaptation on a Humanoid iCub Robot Using a Spiking Cerebellar Model. IEEE Trans. Cybern.. 50(11), 4744–57

Nguyen-Vu, T. B., Zhao, G. Q., Lahiri, S., Kimpo, R. R., Lee, H., Ganguli, S., Shatz, C. J., & Raymond, J. L. (2017). A saturation hypothesis to explain both enhanced and impaired learning with enhanced plasticity. Elife, 6, e20147.

Nobukawa, S., & Nishimura, H. (2016). Chaotic resonance in coupled inferior olive neurons with the Llinás approach neuron model. Neural Comput., 28(11), 2505–32.

Paige, G. (1992). Senescence of human visual-vestibular interactions. 1. Vestibulo-ocular reflex and adaptive plasticity with aging. J. Vestibular Res.: equilibrium & orientation, 2(2), 133–51.

Palay, S. L., & Chan-Palay, V. (2012). Cerebellar cortex: cytology and organization. Springer Science & Business Media.

Pernice, H. F., Schieweck, R., Jafari, M., Straub, T., Bilban, M., Kiebler, M. A., & Popper, B. (2019). Altered Glutamate Receptor Ionotropic Delta Subunit 2 Expression in Stau2-Deficient Cerebellar Purkinje Cells in the Adult Brain. Int. J. Molecular Sciences, 20(7), 1797.

Peterka, R., Black, F., & Schoenhoff, M. (1990). Age-related changes in human vestibulo-ocular reflexes: sinusoidal rotation and caloric tests. J. Vestib. Res., 1(1), 49–59.

Piirtola, M., & Era, P. (2006). Force platform measurements as predictors of falls among older people–a review. Gerontology, 52(1), 1–16.

Renovell, A., Giner, J., & Portoles, M. (2001). Loss of granule neurons in the aging human cerebellar cortex. Int. J. Developmental Biol., 40(S1), S193–S194.

Robinson, D. (1981). The use of control systems analysis in the neurophysiology of eye movements. Annual Rev. Neurosci., 4(1), 463–503.

Ros, E., Carrillo, R. R., Ortigosa, E. M., Barbour, B., & Agís, R. (2006). Event-driven simulation scheme for spiking neural networks using lookup tables to characterize neuronal dynamics. Neural Comput., 18(12), 2959–93.

Roth, A., & Häusser, M. (2001). Compartmental models of rat cerebellar Purkinje cells based on simultaneous somatic and dendritic patch-clamp recordings. J. Physiol., 535, 445–472.

Santina, C. C., Cremer, P. D., Carey, J. P., & Minor, L. B. (2001). The Vestibulo-Ocular Reflex during Self-Generated Head Movements by Human Subjects with Unilateral Vestibular Hypofunction. Annals of the New York Academy of Sciences, 942(1), 465–466.

Sargolzaei, A., Abdelghani, M., Yen, K. K., & Sargolzaei, S. (2016). Sensorimotor control: computing the immediate future from the delayed present. BMC bioinformatics, 17(7), 501–509.

Schweighofer, N., Doya, K., & Kawato, M. (1999). Electrophysiological properties of inferior olive neurons: a compartmental model. J. Neurophysiol., 82(2), 804–817.

Schweighofer, N., Lang, E. J., & Kawato, M. (2013). Role of the olivo-cerebellar complex in motor learning and control. Front Neural Circuits, 7, 94.

Shim, H. G., Jang, D. C., Lee, J., Chung, G., Lee, S., Kim, Y. G., Jeon, D. E., & Kim, S. J. (2017). Long-term depression of intrinsic excitability accompanied by the synaptic depression in the cerebellar Purkinje cells. J. Neurosci., 3464–16.

Shim, H. G., Lee, Y.-S., & Kim, S. J. (2018). The emerging concept of intrinsic plasticity: activity-dependent modulation of intrinsic excitability in cerebellar Purkinje cells and motor learning. Exp. Neurobiol., 27(3), 139.

Skavenski, A. A., & Robinson, D. A. (1973). Role of abducens neurons in vestibuloocular reflex. J. Neurophysiol., 36(4), 724–738.

Smith, P. F. (2016). Age-related neurochemical changes in the vestibular nuclei. Front. Neurol., 7, 20.

Sotelo, C., Llinas, R., & Baker, R. (1974). Structural study of inferior olivary nucleus of the cat: morphological correlates of electrotonic coupling. J. Neurophysiol., 37(3), 541–59.

Sulzer, D., Mosharov, E., Talloczy, Z., Zucca, F. A., Simon, J. D., & Zecca, L. (2008). Neuronal pigmented autophagic vacuoles: lipofuscin, neuromelanin, and ceroid as macroautophagic responses during aging and disease. J. Neurochemistry, 106(1), 24–36.

Tinetti, M. E. (2003). Preventing falls in elderly persons. New England journal of medicine, 348(1), 42–49.

Tokuda, I. T., Hoang, H., Schweighofer, N., & Kawato, M. (2013). Adaptive coupling of inferior olive neurons in cerebellar learning. Neural Networks, 47, 42–50.

Torvik, A., Torp, S., & Lindboe, C. (1986). Atrophy of the cerebellar vermis in ageing: a morphometric and histologic study. J. Neurological Sci., 76(2-3), 283–94.

Turrigiano, G., Abbott, L., & Marder, E. (1994). Activity-dependent changes in the intrinsic properties of cultured neurons. Science, 264(5161), 974–976.

Uusisaari, M., & De Schutter, E. (2011). The mysterious microcircuitry of the cerebellar nuclei. J. Physiol., 589(14), 3441–57.

Viswasom, A. A., Sivan, S., & Jobby, A. (2013). Age related changes in the granule cell number in the human cerebellar cortex. J. Evol. Medical & Dental Sci., 2(16), 2698–2705.

Yamazaki, T., & Tanaka, S. (2005). Neural modeling of an internal clock. Neural Comput., 17(5), 1032–1058.

Yamazaki, T., & Tanaka, S. (2007). The cerebellum as a liquid state machine. Neural Networks, 20, 290–297.

Yamazaki, T., & Tanaka, S. (2009). Computational models of timing mechanisms in the cerebellar granular layer. Cerebellum, 8(4), 423–432.

Yin, D. (1996). Biochemical basis of lipofuscin, ceroid, and age pigment-like fluorophores. Free Radical Biology and Medicine, 21(6), 871–888.

Yuzaki, M. (2013). Cerebellar LTD vs. motor learning—Lessons learned from studying GluD2. Neural Networks, 47, 36–41.

Zalewski, C. K. (2015). Aging of the human vestibular system. Seminars in hearing,

Zanjani, H. S., Vogel, M. W., & Mariani, J. (2016). Deletion of the GluRδ2 receptor in the hotfoot mouse mutant causes granule cell loss, delayed Purkinje cell death, and reductions in Purkinje cell dendritic tree area. Cerebellum, 15(6), 755–66.

Zhang, C., Zhu, Q., & Hua, T. (2010). Aging of cerebellar Purkinje cells. Cell & Tissue Res., 341(3), 341–347.

